# Small-molecule modulation of β-arrestins

**DOI:** 10.1101/2024.12.27.630464

**Authors:** Alem W. Kahsai, Natalia Pakharukova, Henry Y. Kwon, Kunal S. Shah, Jason G. Liang-Lin, Caroline T. del Real, Paul J. Shim, Mason A. Lee, Van A. Ngo, Bowie N. Shreiber, Samuel Liu, Allison M. Schwalb, Emmanuel F. Espinoza, Brittany N. Thomas, Cal A. Kunzle, Jeffrey S. Smith, Jialu Wang, Jihee Kim, Xingdong Zhang, Howard A. Rockman, Alex R. B. Thomsen, Lindsay A.M. Rein, Lei Shi, Seungkirl Ahn, Ali Masoudi, Robert J. Lefkowitz

## Abstract

β-arrestins are multifunctional regulators of G protein-coupled receptor (GPCR) signaling, orchestrating diverse downstream signaling events and physiological responses across the vast GPCR superfamily. While GPCR pharmacology has advanced to target orthosteric and allosteric sites, as well as G proteins and GRKs, comparable chemical tools to study β-arrestins remain lacking. Here, we report the discovery of small-molecule inhibitors that selectively target β-arrestins and delineate their mechanism of action through integrated pharmacological, biochemical, biophysical, and structural analyses. These inhibitors disrupt β-arrestin-engagement with agonist-activated GPCRs, impairing desensitization, internalization, and β-arrestin-dependent functions while sparing G protein–receptor coupling. Cryo-EM, MD simulations, and structure-guided mutagenesis reveal that one modulator, Cmpd-5, engages a cryptic pocket formed by the middle, C-, and lariat loops of β-arrestin1—a critical receptor-binding interface—stabilizing a distinct conformation incompatible with GPCR engagement. Together, these findings provide a mechanistic framework for β-arrestin modulation, introducing transducer-targeted strategies to fine-tune GPCR signaling and guide the development of pathway-specific therapeutics.

## Main

β-arrestins (βarrs) are essential protein communication hubs that regulate ligand activation, signaling, and trafficking of G protein-coupled receptors (GPCRs)^1–4^. GPCRs represent the most prominent family of transmembrane receptors in humans, encompassing over 800 genes, and are the target of approximately one-third of all FDA-approved drugs^5,6^. As versatile adapter protein scaffolds, βarrs play a crucial role in GPCR desensitization, endocytosis, and modulation of signaling pathways^4,7,8^. The mammalian arrestin family includes two widely expressed isoforms, βarr1 and βarr2 (also known as arrestin-2 and -3, respectively), along with two visual arrestins, arrestin-1 and - 4, which are specialized for rod and cone photoreceptors^1,4,8,9^. Upon agonist activation, GPCRs undergo conformational changes that facilitate their interaction with and activation of heterotrimeric G proteins, triggering the production of second messenger molecules and modulation of downstream effectors^1,2^. Subsequent phosphorylation of the agonist-activated receptors by GPCR kinases (GRKs), primarily within the carboxy-terminal tail, enables βarrs to bind to the receptor^1,3,10,11^. This binding sterically hinders further G protein interaction, leading to receptor desensitization, attenuation of G protein-mediated downstream signaling pathways, and receptor internalization via clathrin-coated pits^12–15^.

Beyond their traditional role in signaling termination, βarrs also function as versatile signal transduction units, participating independently or in concert with G proteins in diverse signaling cascades^4,16–18^. βarrs scaffold and facilitate interactions with various signaling mediators, including Mitogen-Activated Protein Kinases (MAPKs), the proto-oncogene kinase Src, Nuclear Factor kappa B (NF-kB), AKT (also known as Protein Kinase B), and Mouse Double Minute 2 Homolog (Mdm2)/p53, among others, as well as various components of the endocytic machinery^4,7,16,19–22^. Their involvement spans many biological processes, encompassing gene expression, metabolism, immune function, motility, and apoptosis^4,23^. Consequently, dysregulated βarr signaling is implicated in a broad spectrum of diseases, including cancer, inflammatory and autoimmune disorders, metabolic syndromes, cardiovascular conditions, and neurological disorders^4,23–29^. Despite their diverse roles, to date, no pharmacological agents directly target βarrs, in contrast to GPCR transducers such as G proteins and GRKs, for which numerous modulators exist^30–36^. As a result, studies of βarrs continue to rely on cumbersome genetic models and animal systems^23,37,38^. These limitations underscore the need for chemical strategies to probe and modulate βarr activity with precision. Here, we report the discovery of the first small molecules that bind and inhibit βarrs, and illuminate their mechanism of action through integrated structural, computational, and functional analyses. Together, these findings introduce new chemical tools that modulate βarrs and establish a mechanistic framework for the selective modulation of βarr-mediated GPCR signaling.

## Results

### Discovery of small-molecule modulators of β-arrestin activity

To identify small molecules capable of modulating βarrs, we implemented a comprehensive, multi-tiered discovery strategy. This began with a differential scanning fluorimetry (DSF)^39,40^ screen to assess compound interactions with purified βarr proteins, followed by functional, biochemical, pharmacological, biophysical and structural analyses including cryo-electron microscopy and molecular dynamics simulations to characterize their binding properties and cellular activity (Fig. 1a and Extended Data Fig. 1). To ensure assay sensitivity and reproducibility, our DSF platform was validated using three established βarr-binding ligands: two highly charged, non-permeant probes–inositol hexakisphosphate (IP_6_), and heparin–and a synthetic phosphopeptide (V_2_Rpp) derived from the C-terminal tail of the vasopressin-2 receptor (V_2_R), a known βarr activator (Extended Data Fig. 1b). Control ligands known to bind βarrs induced marked melting temperature (*T*_m_) shift (Δ*T*_m_) confirming DSF as a robust and sensitive method for detecting small molecule βarr binders (Extended Data Fig. 1b). Based on this validation, a chemically diverse library of 3,500 small molecules (representing over 200K compounds from the National Cancer Institute database) was screened by DSF (Extended Data Fig. 1a, c). Preliminary hits were identified using a threshold (|ΔTₘ| ≥ 1.8 °C), informed by the control ligands (Extended Data Fig. 1a–d). These initial hits were subsequently refined through secondary DSF assays, toxicity filtering and functional screens, yielding 26 modulators with relatively favorable pharmacological profiles (Extended Data Fig. 2a–c). Among these, three chemically and structurally distinct compounds–Cmpd-5, Cmpd-46, and Cmpd-64–were selected for detailed characterization based on inhibitory activity and tractability for further study (Fig. 1b). Schematic overviews and the prioritization cascade are outlined in Extended Data Figs. 1–2.

**Fig. 1:**
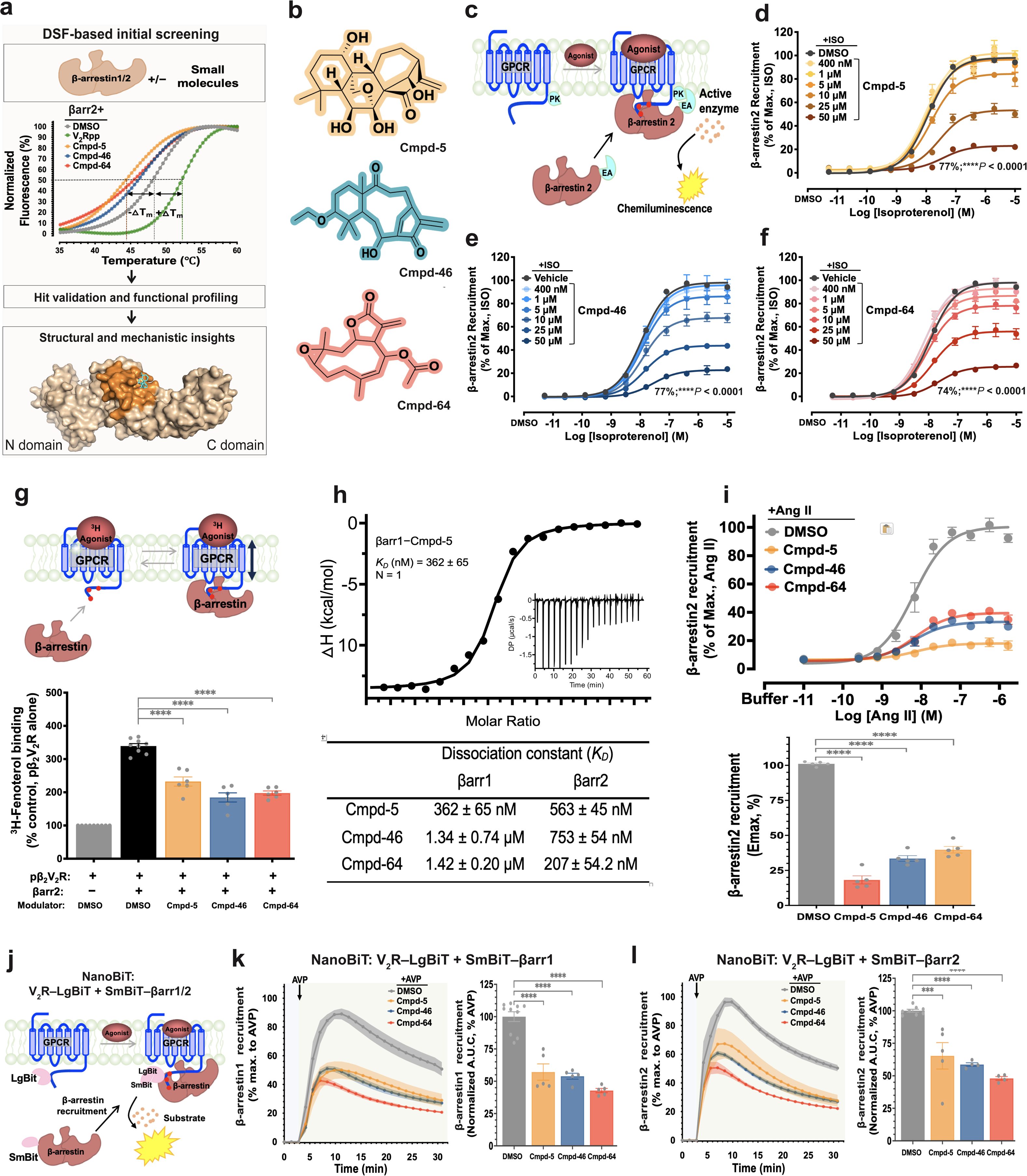
Identification and characterization of small-molecule β-arrestin modulators. **a**, Schematic workflow summarizing the identification and characterization pipeline. DSF melting curves identified βarr binders based on compound-induced Δ*T*_m_ relative to controls. Validated hits underwent functional and mechanistic profiling, including cellular assays, biophysical analyses, MD simulations, and cryo-EM. Molecular surface representation of the βarr1–Cmpd-5 complex, solved by cryo-EM, is shown as a representative structure. **b**, Chemical structures of βarr modulators Cmpd-5, Cmpd-46, and Cmpd-64. **c**, PathHunter β-arrestin recruitment assay. βarr2 tagged with a large β-gal fragment (enzyme acceptor, EA) is co-expressed with β_2_V_2_R fused to a small β-gal donor enzyme fragment (ProLink™; PK) in U2OS cells. Upon receptor activation, βarr2 recruitment restores β-gal activity, generating a chemiluminescent signal. **d**–**f**, Dose-dependent inhibition of βarr2 recruitment by Cmpd-5 (d), Cmpd-46 (e), and Cmpd-64 (f). Compounds were pre-incubated for 30 min before ISO stimulation, reducing E_max_ by 77.0 ± 1.2%, 77.3 ± 1.0%, and 74.4 ± 0.5% at 50 μM, respectively. Data are mean ± s.e.m. (n = 4). **g**, Modulators disrupt βarr2-promoted high-affinity agonist binding. Top, βarr2 association with phosphorylated GPCR stabilizes the active receptor conformation and enhances agonist affinity. Bottom, radiolabeled agonist binding ([^3^H]-Fen, 6 nM) provides a quantitative readout of βarr2 engagement with pβ_2_V_2_R. βarr2 (2 μM, black bar) increased agonist binding relative to pβ_2_V_2_R alone (grey bar); all three modulators (100 μM) inhibited this effect. See Extended Data Fig. 2c for full hit profiles with βarr1 and βarr2. **h**, ITC analysis of modulator binding to βarr1 and βarr2. Top, representative isotherm and raw injection heats (inset) for βarr1 with Cmpd-5 from one of three independent experiments, fit to a one-site model. Bottom, *K_D_* values (mean ± s.e.m.) for all compounds with βarr1 or βarr2. Full thermograms and parameters are provided in Extended Data Fig. 3. **i**, PathHunter β-arrestin recruitment assay at AT_1_R. CHO-K1 cells expressing ProLink-tagged AT_1_R and EA-tagged βarr2 were treated with modulators (50 μM, 30 min) or DMSO, followed by AngII stimulation (1 h). Luminescence was measured as in c. Bottom, Emax values normalized to vehicle (∼100%). **j–l**, NanoBiT luciferase assay of βarr recruitment to V_2_R. **j**, Schematic of the NanoBiT assay with SmBiT–βarr1/2 and LgBiT–V_2_R co-expressed in βarr1/2-knockout HEK293 cells. **k**, **l**, Luminescence measured after modulator pretreatment (50 μM) and AVP stimulation (100 nM). Right, recruitment kinetics and area under the curve (A.U.C) quantification for βarr1 (k) and βarr2 (l). Red dots denote the phosphorylated C-terminus of the receptor (c, g, and j). Data are mean ± s.e.m., normalized to vehicle. One-way ANOVA with Dunnett’s post hoc analysis; \*\*\**P* < 0.001; \*\*\*\**P* < 0.0001.

To assess the functional impact of the candidate modulators, we initially used the PathHunter™ assay with β_2_V_2_R, a chimeric receptor combining β_2_-adrenergic receptor (β₂AR) and the V_2_R C-terminal tail to drive robust βarr2 engagement^41^ (Fig. 1c). This assay uses β-galactosidase (β-gal)-based enzyme fragment complementation (EFC), where agonist-induced βarr2 recruitment brings together β-gal fragments, fused to βarr2 and β_2_V_2_R, reconstituting functional β-gal and producing a chemiluminescent signal proportional to βarr2 recruitment. In an initial single-dose screen, all 26 candidate modulators showed marked inhibition of βarr2 recruitment (Extended Data Fig. 2b). We next prioritized three modulators which showed robust dose-dependent inhibition of isoproterenol (ISO)-induced βarr2 recruitment, with maximal reductions of 77.0% (Cmpd-5), 77.3% (Cmpd-46), and 74.4% (Cmpd-64) at 50 μM (Fig. 1d-f). To complement βarr recruitment assays, we also pharmacologically assessed the impact of candidate modulators on the βarr1/2-promoted high-affinity agonist-bound receptor state using a radioligand binding approach (Fig. 1g). This method quantifies βarr engagement with phosphorylated receptors, here pβ_2_V_2_R, by tracking binding of the radiolabeled agonist ([^3^H]-4-methoxyfenoterol, ^3^H-Fen), which increases reciprocally with βarr coupling^41–43^. Here, agonist binding stabilizes the active receptor conformation, further enhancing βarr engagement. A single-dose screen of all 26 candidates revealed that most modulators reduced βarr-promoted stabilization of the high-affinity state by ≥30% for at least one βarr-isoform (Extended Data Fig. 2c). Notably, the three prioritized compounds each reduced βarr-mediated high-affinity agonist binding by ∼40%, consistent with their effects in the βarr2 recruitment assay (Extended Data Fig. 2c).

To further explore direct interactions, we assessed binding affinities of the modulators to βarr1/2 using isothermal titration calorimetry (ITC), which revealed dissociation constants (*K_D_*) ranging from high nanomolar to low micromolar (Fig. 1h and Extended Data Fig. 3). Cmpd-5 was found not to be isoform-selective, whereas Cmpd-46 showed slight βarr2 preference. In contrast, Cmpd-64 exhibited ∼6-fold selectivity for βarr2 over βarr1. Having confirmed direct binding, we next examined the target specificity of the βarr modulators. Pharmacologically, using a ^3^H-Fen-binding assay, we assessed off-target effects on the Gs heterotrimer (Gαs–βγ), the canonical transducer counterpart of βarrs. None of the modulators affected Gαs–βγ coupling to β₂AR or its ability to stabilize the high-affinity agonist-bound state, as shown at the highest concentration tested (100 μM), indicating a strong transducer selectivity (Extended Data Fig. 4a). To corroborate these findings, we used DSF-based assay to assess direct binding to the Gαs α-subunit and three kinases, common downstream effectors, revealing negligible binding to Gαs, ERK2, c-Src, or p38α (Extended Data Fig. 4b–c). Collectively, these findings confirm that the βarr modulators selectively engage βarrs while sparing the Gs heterotrimer, supporting their targeted action on βarr function.

We next assessed the effect of the three candidate βarr modulators on βarr recruitment at a different receptor system, the angiotensin II type 1 receptor (AT1R), using the same PathHunter™ βarr assay (Fig. 1i). All three compounds suppressed Ang-II–stimulated βarr2 recruitment, consistent with broad inhibition of βarr2 function across distinct receptor engagements. Given their high sequence similarity (∼78%) and functionally divergent roles in certain contexts^4,23^, we next evaluated isoform-specific effects using the NanoBiT βarr assay at V_2_R (Fig. 1j)^44,45^. This split-luciferase complementation system enables real-time monitoring of βarr1 or βarr2 recruitment by reconstituting NanoLuc activity upon agonist (arginine vasopressin, AVP)-stimulated interaction with V_2_R. SmBiT–βarr1 or SmBiT–βarr2 and LgBiT–V_2_R were co-expressed in CRISPR-based βarr1/2-deficient HEK293 cells. Luminescence measured after modulator pretreatment and AVP stimulation confirmed that all three modulators consistently inhibited βarr1/2 recruitment, with similar effects on both isoforms (Fig. 1k,l). Notably, these results are consistent with [^3^H]-Fen binding data showing reduced βarr1/2 coupling to β_2_V_2_R and ITC measurements confirming direct modulator interaction with both βarr isoforms (Fig. 1h, Extended Data Fig. 2c and 3), further supporting the inhibitory effect of the modulators on βarr engagement with agonist-stimulated receptors. Together, these findings demonstrate that the modulators bind βarrs and attenuate agonist-induced GPCR–βarr interactions, indicating broad-spectrum modulation of βarr activities.

### β-arrestin modulators inhibit agonist-promoted receptor desensitization and internalization

Prolonged GPCR activation attenuates agonist response, partly due to βarr binding to phosphorylated receptors, which uncouples them from G proteins and promotes internalization^3,4,12^. To examine how βarr modulators influence this process, we used live-cell biosensors to monitor real-time second messenger dynamics, focusing on cAMP and Ca^2+^. For cAMP, we employed a FRET-based biosensor (ICUE2) in HEK293 cells, stably expressing a fusion of cyan and yellow fluorescent proteins (CFP-Epac2-YFP)^46^ (Fig. 2a). Cells were pretreated with βarr modulators before ISO stimulation of Gαs-coupled β_2_AR, and cAMP levels were measured. Pretreatment with βarr modulators inhibited β_2_AR desensitization, sustaining cAMP signaling and enhancing agonist-induced cAMP accumulation compared to DMSO control (100%) (Fig. 2b,c). To determine whether this effect extended to a Gαq-coupled receptor, we examined Ang-II-induced calcium mobilization in HEK293 cells stably expressing AT_1_R (Fig. 2d)^47^. Consistent with our β_2_AR findings, modulators significantly inhibited AT_1_R desensitization, enhancing Gαq-mediated Ca^2+^ signaling (Fig. 2e,f), leading to an increase in Ca^2+^ mobilization relative to the DMSO control. To confirm that these effects were βarr-dependent, we performed Ca^2+^ signaling assays in parental HEK293 cells and CRISPR/Cas9-generated βarr1/2 knockouts, each transiently expressing AT_1_R (Fig. 2g,h). In parental cells, modulators attenuated desensitization, increasing Ang-II-stimulated Ca^2+^ mobilization by 4- to 5-fold (Fig. 2g). In contrast, βarr1/2-knockout cells showed no further modulation upon treatment, with Ang-II-stimulated Ca^2+^ responses remaining comparable to the untreated control (Fig. 2h). These findings demonstrate that βarrs are essential for modulator-driven inhibition of receptor desensitization, as their absence abolished the effect. Collectively, these results indicate that the modulators impair βarr coupling to GPCRs, thereby preventing receptor desensitization.

**Fig. 2:**
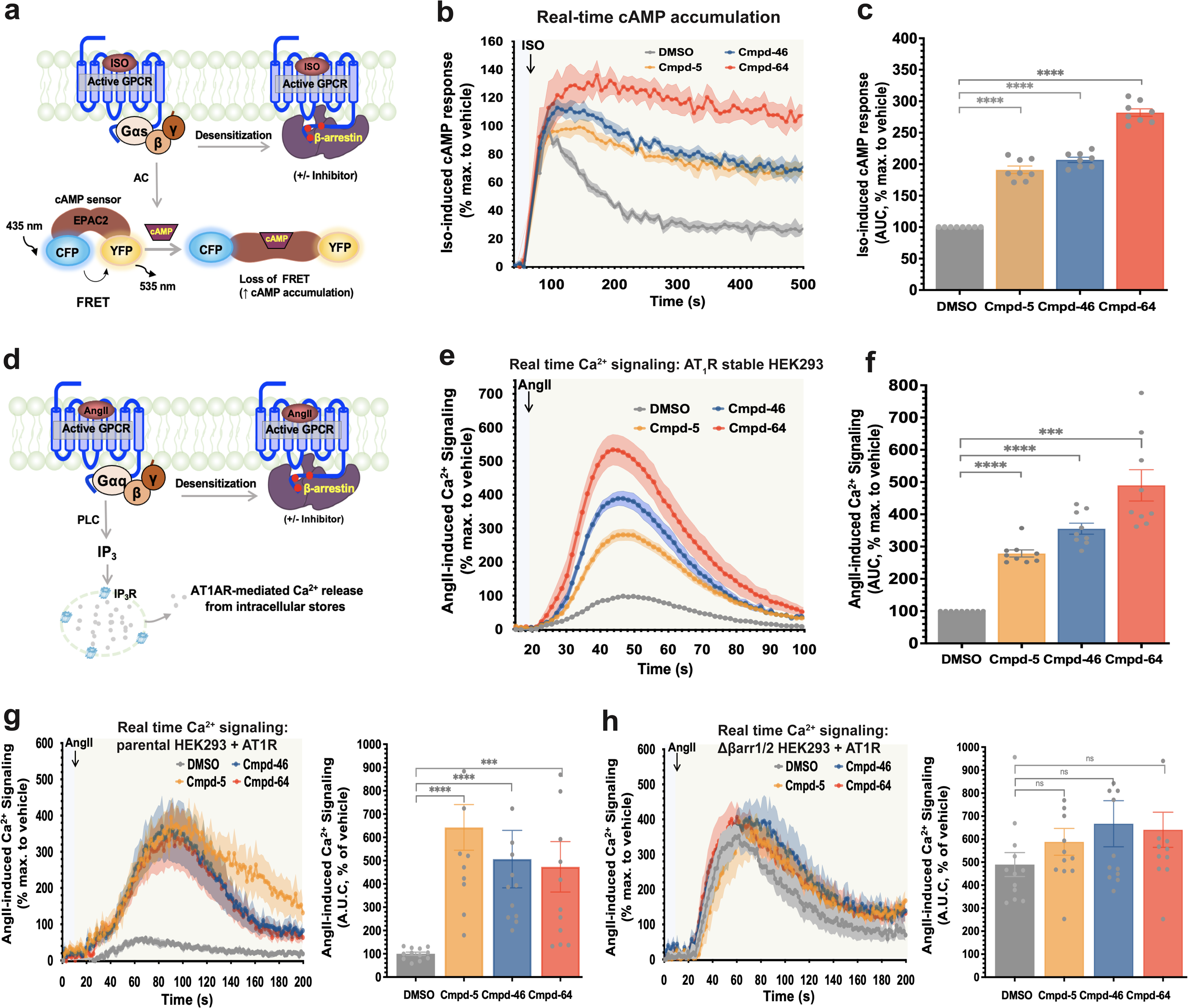
β-arrestin modulators reduce agonist-induced receptor desensitization. **a**, Schematic of the real-time EPAC2-FRET cAMP biosensor (CFP-Epac2-YFP). Upon cAMP binding, increased CFP–YFP distance reduces FRET efficiency. **b**, Kinetic profiles from HEK293 cells stably expressing β₂AR and the cAMP sensor. Cells were pretreated with modulators (40 μM, 5 min) before ISO (10 μM) stimulation. **c**, AUC quantification of FRET responses. βarr modulators increased ISO-induced cAMP accumulation to 191% (Cmpd-5), 207% (Cmpd-46), and 282% (Cmpd-64), relative to DMSO-treated cells (set to 100%) **d**, Schematic of AT_1_R -mediated calcium mobilization via the Gαq pathway. **e**, Real-time Ca^2+^ signaling in HEK293 cells stably expressing AT_1_R. Cells were pretreated with modulators (10 μM, 30 min) before AngII stimulation (30 pM). **F** AUC quantification of Ca^2+^ responses. βarr modulators enhanced AngII-induced Ca^2+^ mobilization by 279% (Cmpd-5), 356% (Cmpd-46), and 490% (Cmpd-64), relative to DMSO controls (100%). **g**, **h**, Real-time Ca^2+^ mobilization in HEK293 cells transiently expressing AT_1_R. g, Parental cells; h, CRISPR βarr1/2-knockout cells. Cells were treated with modulators (10 μM, 30 min) before AngII (120 pM). Right panels in g and h, AUC quantification of Ca^2+^ responses, normalized to AngII alone (100%). Data are mean ± s.e.m. (n = 8). One-way ANOVA with Dunnett’s test; ns, not significant; \*\*\**P* < 0.001; \*\*\*\**P* < 0.0001.

Agonist-induced βarr recruitment to GPCRs facilitates receptor internalization^12,48,49^. Given that all three modulators impaired βarr engagement and receptor desensitization, we next assessed their impact on receptor endocytosis. Specifically, we examined agonist-induced internalization of β_2_V_2_R–βarr2 using the PathHunter β-gal complementation assay in U2OS cells, which stably express β-gal fragments targeted to βarr2 and endosomes (Fig. 3a). Modulator treatment significantly reduced ISO-induced receptor/βarr2 internalization in a dose-dependent manner, with maximal inhibition reaching 93% for Cmpd-5, 88% for Cmpd-46, and 87% for Cmpd-64 at 25 μM (Fig. 3b-d). To independently validate these findings using an orthogonal approach, we employed a bystander bioluminescence resonance energy transfer (BRET) assay to quantify internalization of V₂R–RlucII^50^ (Fig. 3e). This assay measures GPCR trafficking to early endosomes via association with the early endosomal marker 2xFYVE-mVenus. AVP stimulation led to a concentration-dependent increase in the net BRET ratio, indicating V2R endocytosis (Fig. 3f,g). As expected, treatment with all βarr modulators significantly reduced the AVP-induced signal, confirming that internalization to early endosomes, a βarr-dependent process, was inhibited (Fig. 3g). Collectively, these results indicate that the modulators directly bind βarrs, disrupting receptor–βarr interactions and altering this critical function.

**Fig. 3:**
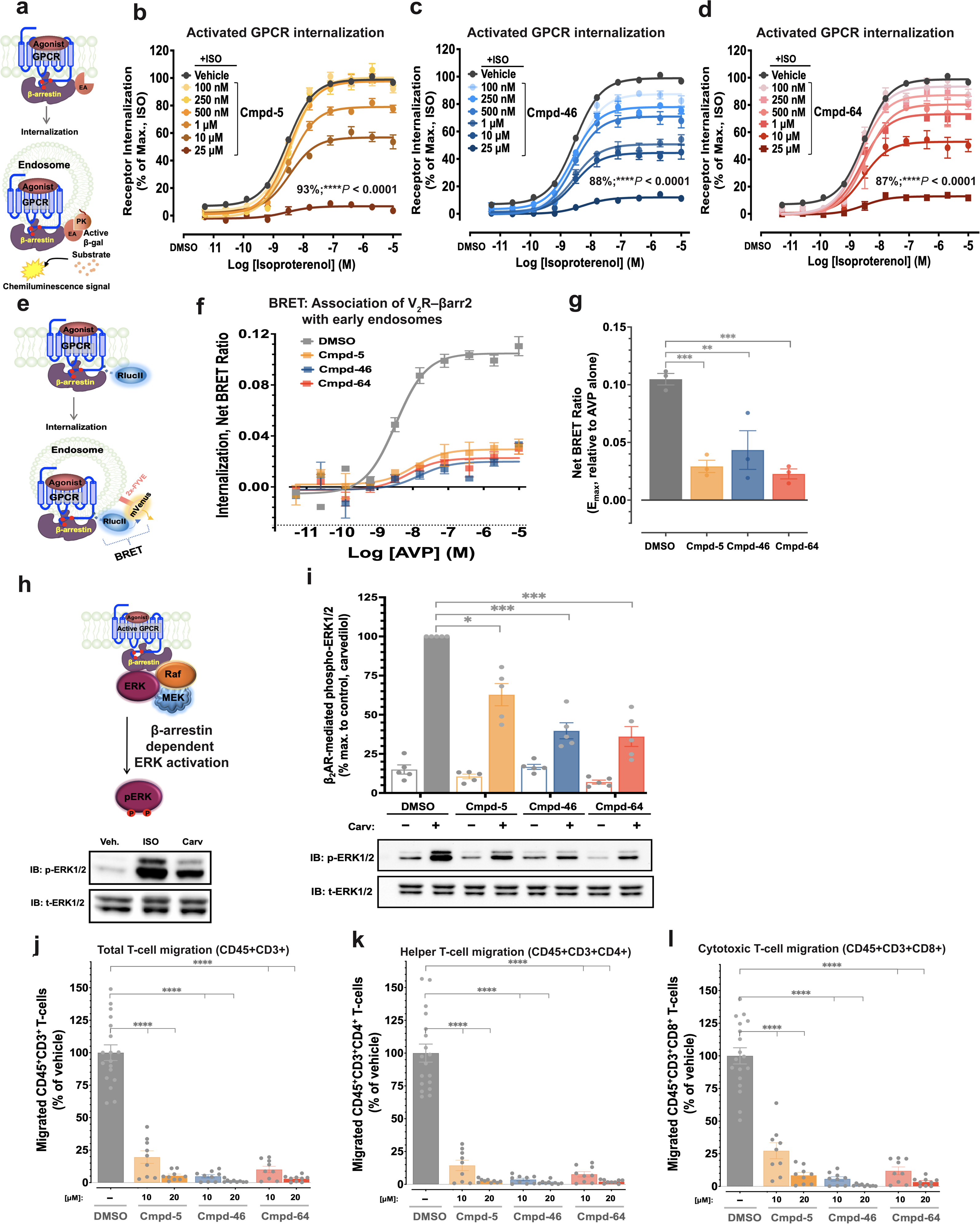
β-arrestin modulators inhibit receptor internalization, downstream signaling, and T cell migration. **a**, Schematic of the PathHunter internalization assay. Agonist-induced βarr2 (EA tagged) recruitment drives internalization of untagged GPCRs into PK-tagged endosomes, reconstituting β-gal activity and generating luminescence. **b**– **d**, Modulators dose-dependently inhibit ISO-induced β_2_V_2_R internalization. Cells were pretreated with modulators for 30 min before ISO stimulation. Cmpd-5 (b), Cmpd-46 (c), and Cmpd-64 (d) reduced internalization, with Emax reductions of 93.2 ± 1.0%, 88.0 ± 0.8%, and 87.1 ± 0.8%, respectively, at 25 μM. **e**, Schematic of BRET-based endocytosis assay. Agonist (AVP) stimulates V_2_R–RlucII internalization and association with the endosomal marker 2xFYVE–mVenus. **f**, Modulators (40 μM, 30 min) reduce AVP-induced increases in net BRET ratio, indicating inhibition of receptor–endosome association. **g**, AUC quantification of BRET responses normalized to AVP alone. **h**, Top, schematic model of βarr-dependent ERK activation, where βarr scaffolds phosphorylated GPCRs to RAF–MEK–ERK signaling components^51,53^. Bottom, Western blot analysis of ERK1/2 phosphorylation in HEK293 cells expressing β_2_AR and stimulated with ISO or carvedilol (10 μM, 5 min). **i**, Modulators (30 μM) attenuate βarr-dependent ERK1/2 phosphorylation in β2AR-expressing HEK293 cells stimulated with carvedilol (10 μM, 5 min). Top, pERK1/2 quantification (relative to Carv alone); bottom, representative blots. **j**–**l**, Modulators impair chemokine-induced T cell migration. Mouse lymphocytes were pretreated with modulators (10 or 20 μM, 30 min) before migration toward CCL19 (100 nM, 2 h). Migrated populations: j, total CD3^+^ T cells (CD45^+^CD3^+^); k, CD4^+^ T helper cells; l, CD8^+^ cytotoxic T cells. Data are mean ± s.e.m. One-way ANOVA (b, g, i–l) and two-way ANOVA (f) with Dunnett’s test. \**P* ≤ 0.05; \*\*\**P* < 0.001; \*\*\*\**P* < 0.0001.

### β-arrestin modulators disrupt βarr-mediated signaling and cellular responses

GPCR-stimulated ERK1/2 activation can occur through a βarr-dependent mechanism ^4,51–53^. To assess the impact of modulators on this pathway, we examined βarr-mediated ERK activation in response to carvedilol, a βarr-biased agonist that suppresses Gαs activation through β_1_/β_2_ARs while inducing moderate βarr-dependent ERK phosphorylation^37,51,54,55^ (Fig. 3h). As anticipated, treatment with Cmpd-5, -46, and -64 attenuated βarr1/2-dependent ERK activation downstream of β_2_AR (Fig. 3i). Furthermore, to validate specificity in the context of ERK activation, we tested whether the modulators influenced EGF-stimulated ERK phosphorylation. Notably, pretreatment with Cmpd-5, -46, and -64 had no effect on EGF-induced ERK activation (Extended Data Fig. 4d), indicating that the modulators do not broadly suppress MAPK signaling. Together with other specificity readouts–including βarr-mediated Gαs coupling, interaction screens with Gαs and kinases, and desensitization analysis in βarr1/2-deficient cells (Fig. 2h; Extended Data Fig. 4a-c)–these results support the conclusion that the modulators selectively target βarr functions downstream of GPCR activation. It is worth noting that although the compounds bind βarrs with measurable affinity in biophysical assays, higher concentrations were needed to elicit cellular effects, possibly due to limited permeability or intracellular stability. These properties may be improved with further chemical optimization.

βarrs scaffold multiple proteins that regulate cell polarity and guide cellular migration downstream of GPCRs, including chemokine receptors^56–58^. To examine whether Cmpd-5, -46, and -64 affect this βarr-dependent function, we assessed their impact on chemotaxis in wild-type mouse T cells (Fig. 3j-l; Extended Data Fig. 5). We focused on CCR7, a chemokine receptor widely expressed across T cell populations and essential for T cell migration^59^. All three βarr modulators markedly disrupted T cell migration, affecting total, helper (CD4^+^), and cytotoxic (CD8^+^) T cell populations (Fig. 3j–l; Extended Data Fig. 5). The extent of inhibition was comparable across cell subsets, with average reductions in chemotaxis of approximately 95%, 99%, and 97% for total, helper, and cytotoxic T cells, respectively, at 20 μM of each compound.

### Structural and molecular mechanism of β-arrestin inhibition

To elucidate the molecular basis of βarr-inhibition by modulators, we determined the cryo-EM structure of the βarr1–Cmpd-5 complex, using Cmpd-5 as a representative modulator for this structural analysis. We prioritized Cmpd-5 based on its ability to bind both βarr isoforms with similar affinity (Fig. 1h, Extended Data Fig. 3) and its favorable aqueous stability, which supported reliable cryo-EM reconstruction. To improve particle alignment in cryo-EM reconstructions, we inserted a BRIL domain at the hinge region of βarr1 and employed an anti-BRIL Fab, BAG2^60^, along with an anti-Fab nanobody (aFabNb)^61^ as fiducial markers (Fig. 4a, Extended Data Fig. 6a). Notably, BRIL insertion did not disrupt βarr1 function, as the modified construct retained the ability to bind Cmpd-5 and V_2_Rpp while stabilizing the high-affinity agonist state of pβ_2_V_2_R (Extended Data Figs. 6b, c), confirming that βarr1-BRIL remained structurally and functionally intact. This approach increased complex stability, aiding data processing, 3D classification, and reconstruction (Extended Data Table 1, Extended Data Fig. 7). Our cryo-EM analysis resolved the βarr1– Cmpd-5 structure at 3.47 Å, revealing well-defined densities for both βarr1 and the modulator Cmpd-5 (Fig. 4a, Extended Data Fig. 8a). The density corresponding to Cmpd-5 is positioned within a distinct pocket in the central crest region of βarr1, bridging the N and C domains. This pocket is formed by the middle loop (residues 129–140), the C-loop (residues 241–249, including β-strand XVII), and the lariat loop (residues 274–300), collectively referred to herein as the MCL site (Fig. 4a–d). Unlike the N-domain groove, which canonically serves as the primary GPCR C-tail binding site^62^, the MCL site appears as a structurally distinct pocket where Cmpd-5 binds, suggesting a potential allosteric role in βarr1 regulation.

**Fig. 4:**
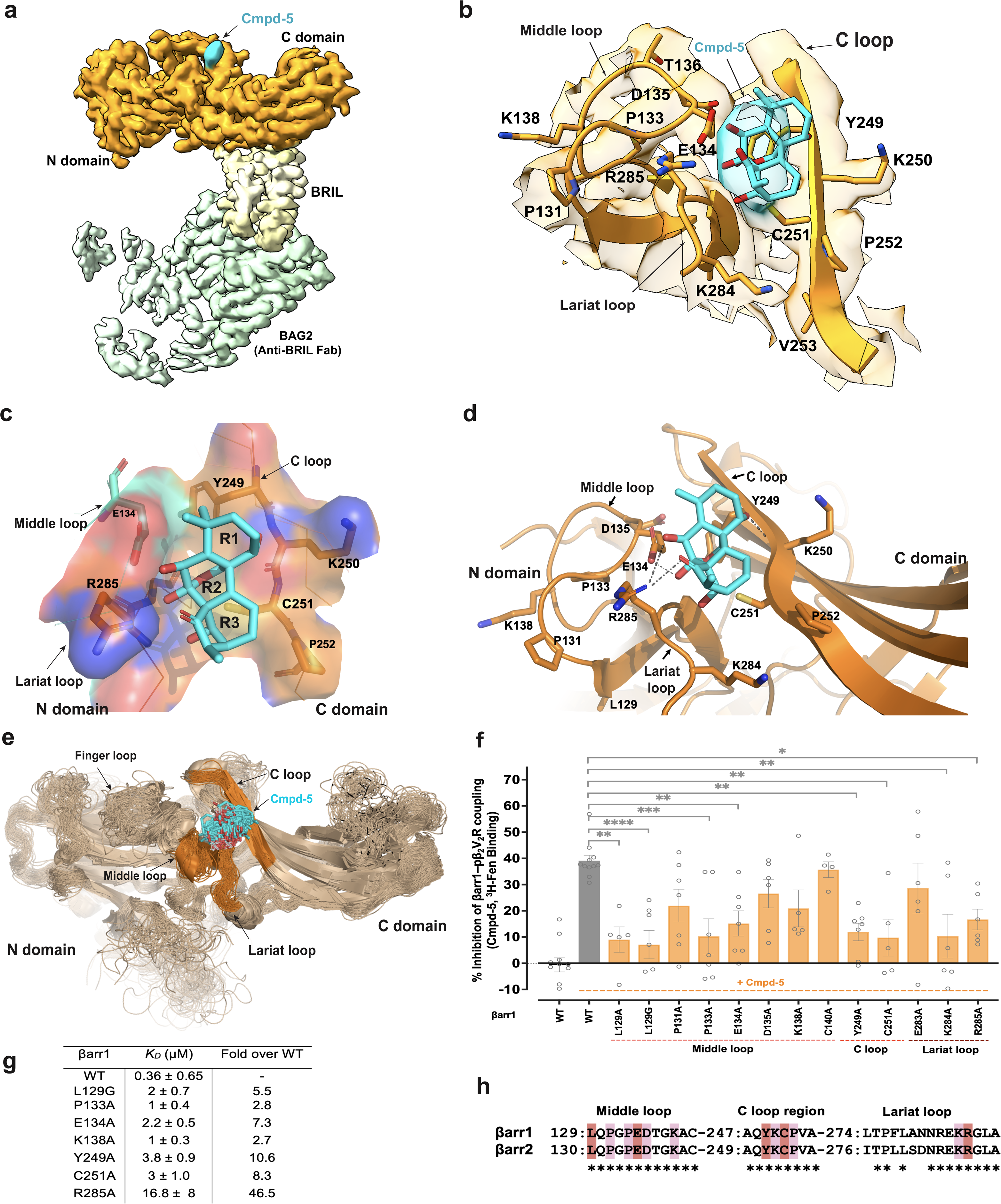
Cmpd-5 binds the MCL cleft in βarr1 and engages a structurally dynamic interface. **a**, Cryo-EM map of βarr1 bound to Cmpd-5 (cyan) showing the N-domain, C-domain, BRIL (pale yellow), and anti-BRIL Fab (BAG2, light green). Map was post-processed with DeepEMhancer and contour level is 0.07. **b**, Cryo-EM close-up of the MCL cleft, showing Cmpd-5 positioned among the middle loop, C-loop, and lariat loop. Key residues are shown as sticks. Map contour level: 0.07. **c**, Electrostatic surface representation of the binding pocket. Cmpd-5 is shown with three moieties: dimethylcyclohexanol (R1), oxabicyclo[2.2.2]octane-diol (R2), and 8-hydroxy-7-methylenebicyclo[3.2.1]octan-6-one (R3). **d**, Cmpd-5 interactions with βarr1, showing probable hydrogen bonds (dashed lines) inferred from structural data, MD simulations, and mutagenesis. **e**, MD simulation snapshots illustrating Cmpd-5 binding within its pocket while accommodating conformational flexibility both in its structure and in the surrounding middle loop, lariat loop, C-loop, and finger loop. **f**, Functional impact of MCL cleft residue mutations on βarr1 modulation, assessed using [^3^H]-Fen binding assays. Cmpd-5 inhibited βarr1-mediated stabilization of the high-affinity β_2_V_2_R state by ∼40%. Data are mean ± s.e.m. from at least 4 experiments (One-way ANOVA with Dunnett’s test. \**P* ≤ 0.05; \*\**P* < 0.01; \*\*\**P* < 0.001; \*\*\*\**P* < 0.0001.). **g**, ITC analysis of Cmpd-5 binding to wild-type and mutant βarr1. *K*_D_ values, mean ± s.e.m., were derived from fits to a one-site binding model. Full representative thermograms and parameters are in Extended Data Fig. 9c–j. **h**, Sequence alignment of rat βarr1 and βarr2 MCL cleft residues. Key residues involved in Cmpd-5 binding are highlighted in red (direct) and pink (indirect).

To validate the binding of Cmpd-5 within the MCL site, we determined the cryo-EM structure of apo βarr1–BRIL at 3.52 Å resolution (Extended Data Fig. 8b, left). In the apo-state map, the absence of density near the MCL site, in contrast to the well-defined density observed in the Cmpd-5-bound βarr1 complex, provided clear evidence for Cmpd-5 occupancy (Fig. 4a and Extended Data Fig. 8b, right). To further support this observation, we generated a difference density map by subtracting the apo-βarr1 density from the Cmpd-5-bound βarr1 structure^63^ revealing additional density at the MCL cleft and corroborating Cmpd-5 binding (Extended Data Fig. 8c). While the observed density was sufficient to confirm Cmpd-5 binding within the MCL cleft, modeling the compound in a specific conformation remained challenging due to its conformational flexibility and peripheral positioning within the pocket. This flexibility–also evident in the MD simulation described below–suggests that while the cryo-EM density confirms binding, it may not fully reflect the range of conformational states adopted by the compound (Fig. 4e, Extended Data Fig. 8d, Extended Data Table 2). The model shows Cmpd-5 engaging both hydrophilic and hydrophobic residues at the βarr1 interface (Fig. 4b–d). Its cyclic hydrocarbon rings align with hydrophobic patches in the C-loop, particularly near Y249 and C251, further stabilizing the interaction. Additionally, E134 (middle loop), K250 (C-loop), and possibly R285 (lariat loop) may mediate polar interactions by engaging the oxygen atoms of Cmpd-5. Although side-chain density for R285 was unresolved, likely due to flexibility, its position suggests accommodation of hydroxyl groups of the compound. These hydrophobic and polar interactions appear critical for stabilizing Cmpd-5 within the MCL site, implying that binding dynamics may extend beyond what static cryo-EM captures (Figs. 4d, e; Extended Data Fig. 8d). To further investigate this, we performed MD simulations to gain deeper insight into Cmpd-5 engagement and its allosteric inhibition of βarr1 over time (Fig. 4e, Extended Data Fig. 8d; Supplementary Video 1 and 2). Consistent with cryo-EM data, simulations showed Cmpd-5 remains stably positioned within the MCL cleft, adopting stable residency in the pocket despite the flexibility of surrounding loops (Supplementary Video 1). The compound also formed polar interactions with R285 and hydrophobic contacts with Y249, C251, and L129 (Supplementary Video 2).

Next, to validate the structural basis of Cmpd-5 binding and understand its inhibitory mechanism, we performed targeted mutagenesis. Leveraging insights from our cryo-EM analysis, we introduced point mutations at key residues within the Cmpd-5 binding site (Extended Data Fig. 9a). We assessed their impact by measuring Cmpd-5–mediated inhibition of βarr1 coupling to pβ_2_V_2_R using [^3^H]-Fen, alongside ITC-based measurements of binding affinities for wild-type and mutant βarr1 variants (Fig. 4f,g; Extended Data Fig. 9b–j). Mutations at L129G, P133A, E134A, Y249A, C251A, K284A, and R285A reduced Cmpd-5 inhibition of βarr1 coupling by 2.4- to 5.5-fold relative to wild-type (Fig. 4f; Extended Data Fig. 9b). In contrast, P131A, D135A, K138A, C140A, and E283A had negligible effects. Building on these observations, ITC provided orthogonal validation of Cmpd-5 binding, complementing the βarr1–pβ_2_V_2_R coupling experiments (Fig. 4g; Extended Data Fig. 9c–j). To refine key determinants, we prioritized a subset of mutants for ITC analysis. Mutations at L129, P133, E134, K138, Y249, C251, and R285 reduced binding affinity by 5.5- to 46.5-fold relative to βarr1-WT. Notably, E134A and R285A impaired both binding and activity, underscoring their role in stabilizing polar interactions (Fig. 4f; Extended Data Fig. 9f,j). For instance, R285A caused a 46.5-fold reduction in affinity, with an enthalpy shift (ΔH: −13.4 kcal/mol in wild-type to −1.5 kcal/mol), indicating loss of stabilizing interactions. Similarly, Y249A and C251A weakened affinity, likely due to disrupted hydrophobic and secondary polar interactions (Extended Data Fig. 9h,i). Interestingly, L129 and P133, though not directly contacting Cmpd-5, still influenced binding, suggesting they stabilize a βarr1 conformation favorable for engagement. Together, cryo-EM, MD simulations, and mutagenesis studies show that Cmpd-5 interacts with βarr1 via a network of hydrophobic and polar contacts, with ITC confirming both enthalpic and entropic contributions. Additionally, sequence alignment of the MCL site between βarr1 and βarr2 (*Rattus norvegicus*) revealed conserved residues, suggesting a shared binding mode across the isoforms (Fig. 4h; Extended Data Figs. 2c, 3a; Fig. 1j,k).

### Distinct conformational state underlying β-arrestin modulation by inhibitor

To elucidate the conformational changes underlying the allosteric inhibition of βarr1 by Cmpd-5, we compared the Cmpd-5–bound and apo-state structures (Fig. 5a–c). While the overall architecture remained largely preserved, Cmpd-5 binding induced marked rearrangements in the central crest region, notably accompanied by an 8° interdomain rotation between the N and C domains (Fig. 5a). Interestingly, difference density maps also revealed additional density near the N domain (Extended Data Fig. 8c), likely reflecting this interdomain twist. In addition to this interdomain twist, Cmpd-5 binding induced bending of the middle loop, repositioning its tip toward the Cmpd-5 site and the C-domain, effectively capping the pocket and reducing its volume. This shift also brought the rear-facing bend of the middle loop into closer proximity with the finger loop (Fig. 5b), potentially contributing to its increased flexibility, consistent with the weaker density observed in the Cmpd-5–bound structure. Supporting this, MD simulations showed that, although the finger loop does not directly contact Cmpd-5, it frequently engages with elements of the N-domain groove (Supplementary Video 1). Cmpd-5 binding also promoted a substantial displacement of the C-loop and induced a helix-to-loop transition in the lariat loop (residues 282–285; Fig. 5c).

**Fig 5:**
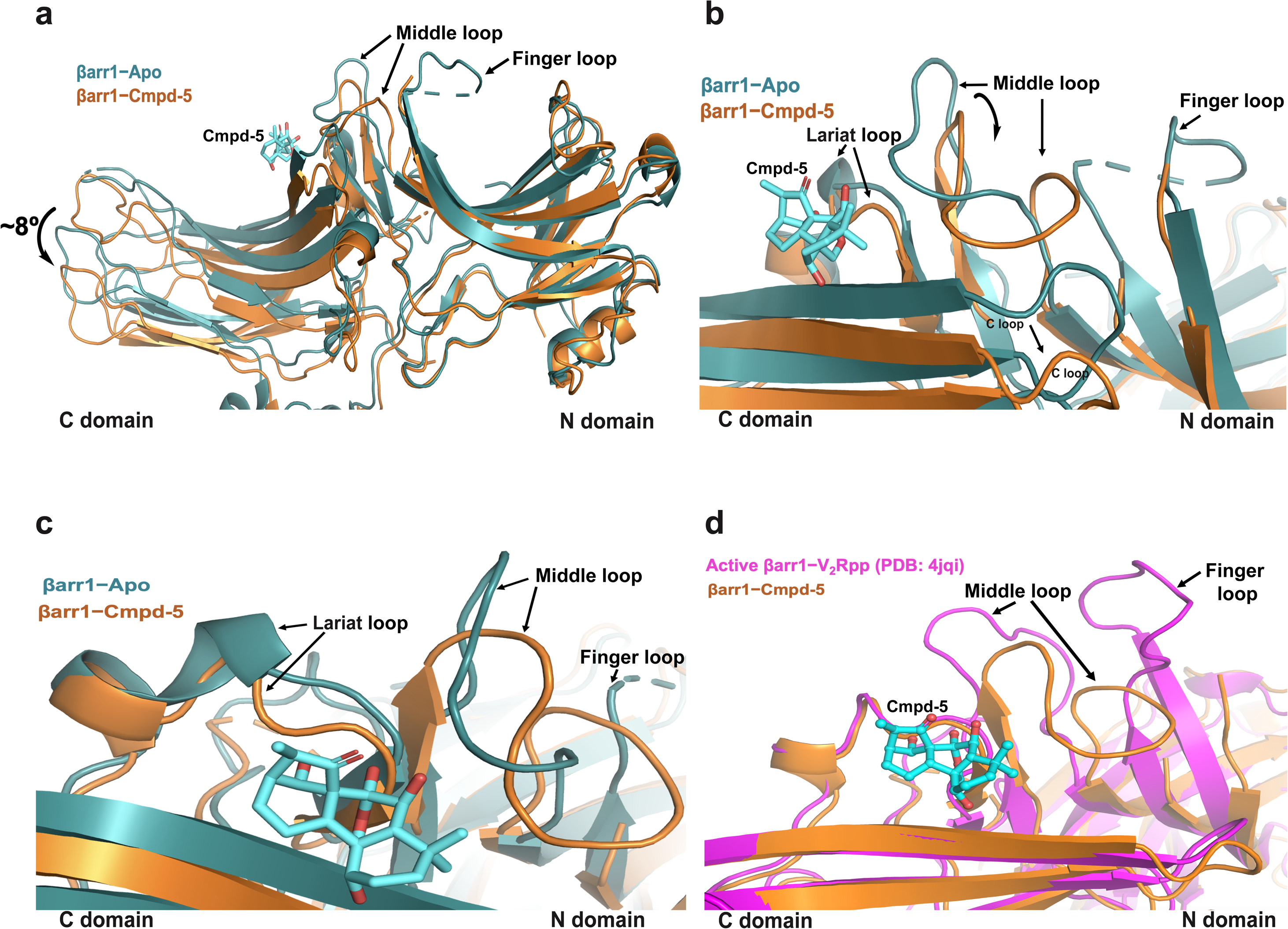
Structural insights into Cmpd-5–mediated allosteric inhibition of βarr1. **a**, Structural comparison of Cmpd-5-bound βarr1 (orange) with apo-βarr1 (deep teal), superimposed by their N domains. Cmpd-5 (cyan) binding induces an 8° interdomain twist, resulting in a poorly resolved finger loop. This twist stabilizes βarr1 in a conformation incompatible with receptor binding or activation. **b**, **c**, Conformational rearrangements in the middle loop, C-loop (b), and lariat loop (c) within the MCL cleft. Cmpd-5 shifts the middle loop toward the N domain and induces a helix-to-loop transition (residues 282– 285) in the lariat loop. Black arrows indicate structural shifts. **d**, Structural overlay of the Cmpd-5–bound βarr1 cryo-EM structure (orange) with the V_2_Rpp-bound βarr1 complex (magenta, PDB: 4JQI), highlighting conformational differences induced by Cmpd-5 binding at the MCL cleft relative to the receptor-bound state.

Superimposing the βarr1–Cmpd-5 structure with the βarr1–V_2_Rpp active-state structure^62^ revealed pronounced differences within the central crest (Fig. 5d). In the active state, the MCL pocket adopts a more open conformation, as the middle loop shifts toward the N domain to accommodate receptor elements such as intracellular loop 2 (ICL2), consistent with multiple GPCR–βarr1 complexes^64–69^. By contrast, Cmpd-5 induces a distinct rearrangement–including bending of the middle loop toward the C-domain, an 8° interdomain twist, and concerted shifts in the lariat and C-loops–that collectively constrict the MCL cleft (Fig. 5a–d; Extended Data Fig. 10a), occluding the binding of elements such as ICL2 and rendering it incompatible with receptor engagement (Extended Data Fig. 10b–d). As a result, the Cmpd-5–bound βarr1 adopts a conformation distinct from both canonical active and inactive states, as also supported by MD simulations (Extended Data Fig. 10e). These structural rearrangements are consistent with an allosteric locking mechanism, in which Cmpd-5 stabilizes a distinct inactive-like βarr1 conformation that prevents receptor engagement, thereby imparing downstream signaling.

## Discussion

βarrs play a central role in regulating diverse GPCR signaling pathways, influencing both physiological functions and pathophysiological processes^4,16,23,38^. Their functions have largely been investigated through genetic approaches, including knockouts, gene silencing, and more recently CRISPR/Cas9-based editing^4,23,37,70–72^. While invaluable, these approaches lack reversibility and temporal precision; in addition, CRISPR/Cas9-mediated knockouts can trigger compensatory mechanisms that obscure the physiological role of βarrs^37,73^. Given these constraints, small-molecule βarr modulators may offer a reversible, precise alternative and serve as tools to probe βarr function, with potential for future preclinical use. Here, we report the discovery of the first small-molecule βarr inhibitors, identified through biophysical screening and characterized via biochemical, pharmacological, functional, structural, and molecular dynamics simulation analyses. Among these, three inhibitors–Cmpd-5, Cmpd-46, and Cmpd-64–were comprehensively characterized as modulators that disrupt βarr interactions with active GPCRs, impairing receptor desensitization, internalization, and downstream βarr-mediated signaling.

Given the pivotal role of βarrs in desensitization^1,3,10,11^, we found that βarr modulators prolong G protein signaling by attenuating receptor desensitization and sustaining second messenger responses via β_2_AR–Gαs and AT_1_R–Gαq coupling.In contrast, these effects were minimal in βarr1/2 knockout cells, indicating selective inhibition of βarrs with limited off-target activity. Similarly, in a radiolabeled β-agonist binding assay, all three modulators inhibited βarr coupling and βarr-promoted high-affinity receptor states without altering Gαs–βγ coupling, the canonical transducer counterpart of βarrs, consistent with an allosteric inhibitory mechanism. Together, these findings support a βarr-directed mode of action that spares G protein function, suggesting favorable specificity. Consistent with this, the modulators also exhibited little to no measurable interaction with the Gαs α-subunit or representative kinases tested, reinforcing a selective profile for βarrs. While encouraging, broader evaluation across diverse signaling contexts will be important in the future to fully define the specificity of these compounds. These results also suggest that these βarr inhibitors may stabilize an inactive-like βarr conformation, as receptor coupling requires an active βarr state, thus biasing GPCR signaling toward the G protein pathway. This indirect shift in signal output highlights the potential for small-molecule βarr modulators to fine-tune GPCR responses. Importantly, most clinically used GPCR-targeting drugs were developed before the recognition of biased signaling and typically engage both G protein- and βarr-mediated pathways in a balanced manner^5,6,74,75^. In contrast, βarr modulators such as those described here may represent a strategy to more selectively disengage βarr signaling, as demonstrated by their effect on desensitization. Indeed, such functional bias could be particularly advantageous in settings where βarr-mediated activities are detrimental. For example, in asthma, βarr-driven attenuation of β_2_AR signaling contributes to airway hyperresponsiveness, whereas G protein signaling promotes bronchodilation^26,76,77^.

Beyond desensitization, βarrs organize GPCR-associated signaling complexes to mediate βarr-dependent G protein-independent pathways, including ERK1/2 activation through the c-Raf–MEK–ERK cascade, acting as scaffolds and allosteric regulators across diverse signaling networks^4,7,16,21,37,51,78^. All three modulators inhibited βarr-dependent ERK activation via β_2_AR, indicating that they interfere with βarr-mediated signaling. Extending these observations, we found that all three compounds also blocked βarr-mediated chemokine-induced T-cell migration–a function where βarrs regulate cell movement by scaffolding proteins involved in cell polarity and actin dynamics downstream of GPCRs^4,58,79^. This is consistent with previous reports showing reduced CCR4-, CXCR3-, and AT_1_R-mediated chemotaxis in βarr-deficient cells^4,56,80,81^. Altogether, these findings position βarr modulators as versatile tools to probe βarr-regulated signaling, while opening avenues to fine-tune GPCR pathways in both physiological and pathophysiological contexts^4,23^.

To understand the structural and mechanistic basis underlying βarr modulation, we employed cryo-EM, MD simulations, and structure-guided mutagenesis, revealing that one of the modulators, Cmpd-5, binds within the MCL cleft–a pocket formed by the middle, C, and lariat loops. Interestingly, this region aligns with the receptor-binding interface in active-state βarr1, where it accommodates intracellular GPCR elements, particularly ICL2^62,64–69^. In the active conformation, the pocket is notably expanded, a result of activation-associated structural changes including an ∼20° interdomain rotation–a hallmark of βarr1 activation. By contrast, Cmpd-5 occupies the same site but induces a distinct set of conformational rearrangements–including bending and reorientation of the middle loop toward the C-domain, C-loop displacement, a helix-to-loop transition in the lariat loop, and, notably, a smaller ∼8° interdomain twist–that together act to constrict the pocket. This region plays a key role in βarr activation and receptor engagement, and such rearrangements may reduce the volume of the cleft and, in doing so, limit accommodation of receptor elements such as ICL2. Collectively, our cryo-EM analysis and MD simulations^82,83^ suggest that Cmpd-5 modulates βarr1 by stabilizing an inactive-like conformation, distinct from both the canonical active and inactive states. Given that βarrs adopt multiple conformations depending on cellular context, the ability of Cmpd-5 to stabilize a distinct inactive-like state advances our understanding of these dynamic signaling proteins^9,84,85^.

Although our structural findings are consistent with an allosteric mechanism, the proximity of the Cmpd-5 binding site to receptor-interacting elements, particularly ICL2, means that partial competitive interference cannot be entirely ruled out. Viewed as a whole, these insights underscore the MCL cleft, a previously uncharacterized regulatory site, as a tractable target for small-molecule modulation of βarr function within GPCR signaling. Furthermore, the intrinsic conformational flexibility of βarrs raises the possibility that future modulators, or optimized analogs based on existing chemical scaffolds, may preferentially stabilize alternative, noncanonical βarr states–offering a path toward more refined pharmacological control of βarrs. Although this study centers on Cmpd-5, it establishes a structural framework for exploring divergent mechanisms of βarr modulation and deepening our understanding of βarr regulation.

In summary, this work uncovers small-molecule modulators of βarrs and delineates the structural basis of βarr1 inhibition by Cmpd-5, which stabilizes a distinct inactive-like conformation by constricting the MCL cleft, a key receptor-binding interface. These findings reveal a previously uncharacterized allosteric site and provide a mechanism by which βarr–GPCR interactions are inhibited via a structurally defined mechanism, offering insight into βarr conformational dynamics and laying a foundation for understanding βarr structure–function relationships. In addition to introducing chemical approaches to probe βarr biology, this study also signals a conceptual shift, moving from traditional receptor-centric strategies to the direct modulation of intracellular transducers such as βarrs, presenting yet another avenue to more precisely regulate and fine-tune GPCR signaling.

## Reporting summary

Further information on research design is available in the Nature Portfolio Reporting Summary linked to this article.

## Data availability

The authors declare that all data supporting the findings of this study are included in the article, Extended Data, Supplementary Information (including Supplementary Tables), or Supplementary Videos. The coordinates and cryo-EM maps have been deposited in the Protein Data Bank (PDB) and the Electron Microscopy Data Bank (EMDB) under accession codes 9DNM and EMD-47042 for βarr1–Cmpd-5, and 9DNG and EMD-47040 for apo βarr1. Additional data and materials, including those supporting the Supplementary Information, are available upon request.

## Code availability

No custom code was used in this study.

## Inclusion & ethics statement

All authors meet the authorship criteria required by *Nature Portfolio journals* and contributed meaningfully to the study. Authorship was determined collaboratively and was not influenced by gender, seniority, or institutional affiliation. Roles were agreed upon in advance, and the work was conducted responsibly and ethically, following institutional standards and inclusive research practices.

## Author contributions

A.W.K. and R.J.L. conceived the study. A.W.K., H.Y.K., K.S.S., J.G.L., C.T.dR., P.J.S., M.A.L., B.N.S., S.L., A.M.S., E.F.E., C.A.K., J.W., and S.A. designed, conducted, and analyzed drug screening, biophysical assays, *in vitro* binding studies, and biochemical and cellular functional assays. A.W.K., N.P., K.S.S., J.G.L., C.T.dR., P.J.S., B.N.S., B.N.T., J.K., and X.Z. contributed to construct design, reagent preparation, protein purification, and quality control. L.A.M.R. and J.S.S. conducted T cell migration assay. N.P., with assistance from B.N.T., prepared cryo-EM samples and performed data acquisition. N.P. performed data processing, model building, and refinement. V.A.G. and L.S. conducted MD simulations and data analysis. A.W.K., N.P., L.S., A.M., and R.J.L. analyzed and interpreted the structural data. A.W.K., H.A.R., A.R.B.T., L.A.M.R., L.S., S.A., and R.J.L coordinated and supervised research. A.W.K. and R.J.L. drafted the original manuscript, and A.W.K., N.P., A.M., and R.J.L. finalized the paper with input from all authors.

## Acknowledgments

R.J.L. is an investigator with the Howard Hughes Medical Institute (HHMI). This work was supported in part by the U.S. National Institutes of Health, National Heart, Lung, and Blood Institute (Grant HL16037 to R.J.L.; Grants T32HL007101 and HL16037-45S1 to A.W.K.; and Grants R35GM147088 and R21CA243052 to A.R.B.T.), the National Institute on Drug Abuse – Intramural Research Program (Z1A DA000606 to L.S.), and the St. Jude Children’s Research Hospital Collaborative Research Consortium on G protein-coupled receptors (GPCRs). We thank the National Cancer Institute (NCI) Developmental Therapeutics Program (DTP) Open Chemical Repository for providing compound libraries and additional compounds for this research. Cryo-EM data were screened and collected using the Krios at Duke University’s Shared Materials Instrumentation Facility (SMIF), a member of the North Carolina Research Triangle Nanotechnology Network (RTNN), supported by the National Science Foundation (award number ECCS-2025064) as part of the National Nanotechnology Coordinated Infrastructure (NNCI). We are grateful to Dr. Nilakshee Bhattacharya at SMIF for assistance with microscope operation and data collection. This work utilized the computational resources of the NIH HPC Biowulf cluster (http://hpc.nih.gov). We also acknowledge the Flow Cytometry Shared Resource at Duke Cancer Institute. We thank Dr. Stéphane Laporte (McGill University, Montreal, Quebec, Canada) for the generous gift of CRISPR/Cas9 βarr1/2-knockout (KO) HEK293 cell lines and Dr. Irving Wainer (Laboratory of Clinical Investigation, National Institute on Aging Intramural Research Program, Baltimore, MD) for providing [^3^H] (R,R′)-4-methoxyfenoterol. N.P. is supported by postdoctoral fellowships from the Human Frontier Science Program (LT000174/2018) and the European Molecular Biology Organization (ALTF 1071-2017). P.J.S. and H.Y.K. are supported by Research Fellowships from the Sarnoff Cardiovascular Research Foundation. Our gratitude also goes to Icee (Yangyang) Li for her diligent secretarial support, as well as to Dr. Li-Yin Huang, Stephanie Martin, Xinrong Jiang, and William Capel for generously sharing their technical expertise.

## References

1. Lefkowitz, R.J. A brief history of G-protein coupled receptors (Nobel Lecture). Angew Chem Int Ed Engl 52, 6366–6378 (2013).

2. Pierce, K.L., Premont, R.T. & Lefkowitz, R.J. Seven-transmembrane receptors. Nat Rev Mol Cell Biol 3, 639–650 (2002).

3. Lohse, M.J., Benovic, J.L., Codina, J., Caron, M.G. & Lefkowitz, R.J. beta-Arrestin: a protein that regulates beta-adrenergic receptor function. Science 248, 1547–1550 (1990).

4. Peterson, Y.K. & Luttrell, L.M. The Diverse Roles of Arrestin Scaffolds in G Protein-Coupled Receptor Signaling. Pharmacol Rev 69, 256–297 (2017).

5. Hauser, A.S., Attwood, M.M., Rask-Andersen, M., Schioth, H.B. & Gloriam, D.E. Trends in GPCR drug discovery: new agents, targets and indications. Nat Rev Drug Discov 16, 829–842 (2017).

6. Sriram, K. & Insel, P.A. G Protein-Coupled Receptors as Targets for Approved Drugs: How Many Targets and How Many Drugs? Mol Pharmacol 93, 251–258 (2018).

7. Kahsai, A.W., et al. Signal transduction at GPCRs: Allosteric activation of the ERK MAPK by beta-arrestin. Proc Natl Acad Sci U S A 120, e2303794120 (2023).

8. Gurevich, V.V. & Gurevich, E.V. Extensive shape shifting underlies functional versatility of arrestins. Curr Opin Cell Biol 27, 1–9 (2014).

9. Kahsai, A.W., Pani, B. & Lefkowitz, R.J. GPCR signaling: conformational activation of arrestins. Cell Res 28, 783–784 (2018).

10. Benovic, J.L., et al. Functional desensitization of the isolated beta-adrenergic receptor by the beta-adrenergic receptor kinase: potential role of an analog of the retinal protein arrestin (48-kDa protein). Proc Natl Acad Sci U S A 84, 8879–8882 (1987).

11. Pitcher, J.A., Freedman, N.J. & Lefkowitz, R.J. G protein-coupled receptor kinases. Annu Rev Biochem 67, 653–692 (1998).

12. Ferguson, S.S. Evolving concepts in G protein-coupled receptor endocytosis: the role in receptor desensitization and signaling. Pharmacol Rev 53, 1–24 (2001).

13. Laporte, S.A., et al. The beta2-adrenergic receptor/betaarrestin complex recruits the clathrin adaptor AP-2 during endocytosis. Proc Natl Acad Sci U S A 96, 3712–3717 (1999).

14. Wolfe, B.L. & Trejo, J. Clathrin-dependent mechanisms of G protein-coupled receptor endocytosis. Traffic 8, 462–470 (2007).

15. Grady, E.F., Bohm, S.K. & Bunnett, N.W. Turning off the signal: mechanisms that attenuate signaling by G protein-coupled receptors. Am J Physiol 273, G586–601 (1997).

16. Lefkowitz, R.J. & Shenoy, S.K. Transduction of receptor signals by beta-arrestins. Science 308, 512–517 (2005).

17. Thomsen, A.R.B., et al. GPCR-G Protein-beta-Arrestin Super-Complex Mediates Sustained G Protein Signaling. Cell 166, 907–919 (2016).

18. Smith, J.S., et al. Noncanonical scaffolding of G(alphai) and beta-arrestin by G protein-coupled receptors. Science 371(2021).

19. Xiao, K., et al. Global phosphorylation analysis of beta-arrestin-mediated signaling downstream of a seven transmembrane receptor (7TMR). Proc Natl Acad Sci U S A 107, 15299–15304 (2010).

20. Xiao, K., et al. Functional specialization of beta-arrestin interactions revealed by proteomic analysis. Proc Natl Acad Sci U S A 104, 12011–12016 (2007).

21. Pakharukova, N., Masoudi, A., Pani, B., Staus, D.P. & Lefkowitz, R.J. Allosteric activation of proto-oncogene kinase Src by GPCR-beta-arrestin complexes. J Biol Chem 295, 16773–16784 (2020).

22. Ahn, S., Shenoy, S.K., Luttrell, L.M. & Lefkowitz, R.J. SnapShot: beta-Arrestin Functions. Cell 182, 1362–1362 e1361 (2020).

23. Wess, J., Oteng, A.B., Rivera-Gonzalez, O., Gurevich, E.V. & Gurevich, V.V. beta-Arrestins: Structure, Function, Physiology, and Pharmacological Perspectives. Pharmacol Rev 75, 854–884 (2023).

24. Freedman, N.J. & Shenoy, S.K. Regulation of inflammation by beta-arrestins: Not just receptor tales. Cell Signal 41, 41–45 (2018).

25. Bagnato, A. & Rosano, L. New Routes in GPCR/beta-Arrestin-Driven Signaling in Cancer Progression and Metastasis. Front Pharmacol 10, 114 (2019).

26. Bond, R.A., Lucero Garcia-Rojas, E.Y., Hegde, A. & Walker, J.K.L. Therapeutic Potential of Targeting ss-Arrestin. Front Pharmacol 10, 124 (2019).

27. Thathiah, A., et al. beta-arrestin 2 regulates Abeta generation and gamma-secretase activity in Alzheimer’s disease. Nat Med 19, 43–49 (2013).

28. Fereshteh, M., et al. beta-Arrestin2 mediates the initiation and progression of myeloid leukemia. Proc Natl Acad Sci U S A 109, 12532–12537 (2012).

29. Rein, L.A., et al. beta-Arrestin2 mediates progression of murine primary myelofibrosis. JCI Insight 2(2017).

30. Schlegel, J.G., et al. Macrocyclic Gq Protein Inhibitors FR900359 and/or YM-254890-Fit for Translation? ACS Pharmacol Transl Sci 4, 888–897 (2021).

31. Schrage, R., et al. The experimental power of FR900359 to study Gq-regulated biological processes. Nat Commun 6, 10156 (2015).

32. Xiong, X.F., et al. Structure-Activity Relationship Studies of the Natural Product G(q/11) Protein Inhibitor YM-254890. ChemMedChem 14, 865–870 (2019).

33. Dai, S.A., et al. State-selective modulation of heterotrimeric Galphas signaling with macrocyclic peptides. Cell 185, 3950–3965 e3925 (2022).

34. Bouley, R.A., et al. A New Paroxetine-Based GRK2 Inhibitor Reduces Internalization of the mu-Opioid Receptor. Mol Pharmacol 97, 392–401 (2020).

35. Thal, D.M., Yeow, R.Y., Schoenau, C., Huber, J. & Tesmer, J.J. Molecular mechanism of selectivity among G protein-coupled receptor kinase 2 inhibitors. Mol Pharmacol 80, 294–303 (2011).

36. Ferrero, K.M. & Koch, W.J. GRK2 in cardiovascular disease and its potential as a therapeutic target. J Mol Cell Cardiol 172, 14–23 (2022).

37. Luttrell, L.M., et al. Manifold roles of beta-arrestins in GPCR signaling elucidated with siRNA and CRISPR/Cas9. Sci Signal 11(2018).

38. Gurevich, V.V. & Gurevich, E.V. Plethora of functions packed into 45 kDa arrestins: biological implications and possible therapeutic strategies. Cell Mol Life Sci 76, 4413–4421 (2019).

39. Niesen, F.H., Berglund, H. & Vedadi, M. The use of differential scanning fluorimetry to detect ligand interactions that promote protein stability. Nat Protoc 2, 2212–2221 (2007).

40. Renaud, J.P., et al. Biophysics in drug discovery: impact, challenges and opportunities. Nat Rev Drug Discov 15, 679–698 (2016).

41. Ahn, S., et al. Allosteric "beta-blocker" isolated from a DNA-encoded small molecule library. Proc Natl Acad Sci U S A 114, 1708–1713 (2017).

42. Gurevich, V.V., Pals-Rylaarsdam, R., Benovic, J.L., Hosey, M.M. & Onorato, J.J. Agonist-receptor-arrestin, an alternative ternary complex with high agonist affinity. J Biol Chem 272, 28849–28852 (1997).

43. Toll, L., et al. Thermodynamics and docking of agonists to the beta(2)-adrenoceptor determined using [(3)H](R,R’)-4-methoxyfenoterol as the marker ligand. Mol Pharmacol 81, 846–854 (2012).

44. Dixon, A.S., et al. NanoLuc Complementation Reporter Optimized for Accurate Measurement of Protein Interactions in Cells. ACS Chem Biol 11, 400–408 (2016).

45. Pandey, S., et al. Intrinsic bias at non-canonical, beta-arrestin-coupled seven transmembrane receptors. Mol Cell 81, 4605–4621 e4611 (2021).

46. Violin, J.D., et al. beta2-adrenergic receptor signaling and desensitization elucidated by quantitative modeling of real time cAMP dynamics. J Biol Chem 283, 2949–2961 (2008).

47. Li, A., Liu, S., Huang, R., Ahn, S. & Lefkowitz, R.J. Loss of biased signaling at a G protein-coupled receptor in overexpressed systems. PLoS One 18, e0283477 (2023).

48. Oakley, R.H., Laporte, S.A., Holt, J.A., Caron, M.G. & Barak, L.S. Differential affinities of visual arrestin, beta arrestin1, and beta arrestin2 for G protein-coupled receptors delineate two major classes of receptors. J Biol Chem 275, 17201–17210 (2000).

49. Laporte, S.A., Oakley, R.H., Holt, J.A., Barak, L.S. & Caron, M.G. The interaction of beta-arrestin with the AP-2 adaptor is required for the clustering of beta 2-adrenergic receptor into clathrin-coated pits. J Biol Chem 275, 23120–23126 (2000).

50. Smith, J.S., et al. C-X-C Motif Chemokine Receptor 3 Splice Variants Differentially Activate Beta-Arrestins to Regulate Downstream Signaling Pathways. Mol Pharmacol 92, 136–150 (2017).

51. Wisler, J.W., et al. A unique mechanism of beta-blocker action: carvedilol stimulates beta-arrestin signaling. Proc Natl Acad Sci U S A 104, 16657–16662 (2007).

52. Luttrell, L.M., et al. Beta-arrestin-dependent formation of beta2 adrenergic receptor-Src protein kinase complexes. Science 283, 655–661 (1999).

53. Noma, T., et al. Beta-arrestin-mediated beta1-adrenergic receptor transactivation of the EGFR confers cardioprotection. J Clin Invest 117, 2445–2458 (2007).

54. Wang, J., et al. Galphai is required for carvedilol-induced beta1 adrenergic receptor beta-arrestin biased signaling. Nat Commun 8, 1706 (2017).

55. Kim, I.M., et al. Beta-blockers alprenolol and carvedilol stimulate beta-arrestin-mediated EGFR transactivation. Proc Natl Acad Sci U S A 105, 14555–14560 (2008).

56. Smith, J.S., et al. Biased agonists of the chemokine receptor CXCR3 differentially control chemotaxis and inflammation. Sci Signal 11(2018).

57. Cheung, R., et al. An arrestin-dependent multi-kinase signaling complex mediates MIP-1beta/CCL4 signaling and chemotaxis of primary human macrophages. J Leukoc Biol 86, 833–845 (2009).

58. McGovern, K.W. & DeFea, K.A. Molecular mechanisms underlying beta-arrestin-dependent chemotaxis and actin-cytoskeletal reorganization. Handb Exp Pharmacol 219, 341–359 (2014).

59. Comerford, I., et al. A myriad of functions and complex regulation of the CCR7/CCL19/CCL21 chemokine axis in the adaptive immune system. Cytokine Growth Factor Rev 24, 269–283 (2013).

60. Mukherjee, S., et al. Synthetic antibodies against BRIL as universal fiducial marks for single-particle cryoEM structure determination of membrane proteins. Nat Commun 11, 1598 (2020).

61. Ereno-Orbea, J., et al. Structural Basis of Enhanced Crystallizability Induced by a Molecular Chaperone for Antibody Antigen-Binding Fragments. J Mol Biol 430, 322–336 (2018).

62. Shukla, A.K., et al. Structure of active beta-arrestin-1 bound to a G-protein-coupled receptor phosphopeptide. Nature 497, 137–141 (2013).

63. Joseph, A.P., et al. Comparing Cryo-EM Reconstructions and Validating Atomic Model Fit Using Difference Maps. J Chem Inf Model 60, 2552–2560 (2020).

64. Bous, J., et al. Structure of the vasopressin hormone-V2 receptor-beta-arrestin1 ternary complex. Sci Adv 8, eabo7761 (2022).

65. Staus, D.P., et al. Structure of the M2 muscarinic receptor-beta-arrestin complex in a lipid nanodisc. Nature 579, 297–302 (2020).

66. Wang, Y., et al. Cryo-EM structure of cannabinoid receptor CB1-beta-arrestin complex. Protein Cell 15, 230–234 (2024).

67. Cao, C., et al. Signaling snapshots of a serotonin receptor activated by the prototypical psychedelic LSD. Neuron 110, 3154–3167 e3157 (2022).

68. Zhou, X.E., et al. Identification of Phosphorylation Codes for Arrestin Recruitment by G Protein-Coupled Receptors. Cell 170, 457–469 e413 (2017).

69. Lee, Y., et al. Molecular basis of beta-arrestin coupling to formoterol-bound beta(1)-adrenoceptor. Nature 583, 862–866 (2020).

70. Kohout, T.A., Lin, F.S., Perry, S.J., Conner, D.A. & Lefkowitz, R.J. beta-Arrestin 1 and 2 differentially regulate heptahelical receptor signaling and trafficking. Proc Natl Acad Sci U S A 98, 1601–1606 (2001).

71. Conner, D.A., et al. beta-Arrestin1 knockout mice appear normal but demonstrate altered cardiac responses to beta-adrenergic stimulation. Circ Res 81, 1021–1026 (1997).

72. Bohn, L.M., et al. Enhanced morphine analgesia in mice lacking beta-arrestin 2. Science 286, 2495–2498 (1999).

73. Lefkowitz, R.J., et al. How carvedilol does not activate beta(2)-adrenoceptors. Nat Commun 14, 7866 (2023).

74. Kahsai, A.W., et al. Multiple ligand-specific conformations of the beta2-adrenergic receptor. Nat Chem Biol 7, 692–700 (2011).

75. Wootten, D., Christopoulos, A., Marti-Solano, M., Babu, M.M. & Sexton, P.M. Mechanisms of signalling and biased agonism in G protein-coupled receptors. Nat Rev Mol Cell Biol 19, 638–653 (2018).

76. Ahn, S., et al. Allosteric modulator potentiates beta2AR agonist-promoted bronchoprotection in asthma models. J Clin Invest 133(2023).

77. Walker, J.K. & DeFea, K.A. Role for beta-arrestin in mediating paradoxical beta2AR and PAR2 signaling in asthma. Curr Opin Pharmacol 16, 142–147 (2014).

78. Luttrell, L.M., et al. Activation and targeting of extracellular signal-regulated kinases by beta-arrestin scaffolds. Proc Natl Acad Sci U S A 98, 2449–2454 (2001).

79. Zoudilova, M., et al. beta-Arrestins scaffold cofilin with chronophin to direct localized actin filament severing and membrane protrusions downstream of protease-activated receptor-2. J Biol Chem 285, 14318–14329 (2010).

80. Hunton, D.L., et al. Beta-arrestin 2-dependent angiotensin II type 1A receptor-mediated pathway of chemotaxis. Mol Pharmacol 67, 1229–1236 (2005).

81. Lin, R., Choi, Y.H., Zidar, D.A. & Walker, J.K.L. beta-Arrestin-2-Dependent Signaling Promotes CCR4-mediated Chemotaxis of Murine T-Helper Type 2 Cells. Am J Respir Cell Mol Biol 58, 745–755 (2018).

82. Han, M., Gurevich, V.V., Vishnivetskiy, S.A., Sigler, P.B. & Schubert, C. Crystal structure of beta-arrestin at 1.9 A: possible mechanism of receptor binding and membrane Translocation. Structure 9, 869–880 (2001).

83. Milano, S.K., Pace, H.C., Kim, Y.M., Brenner, C. & Benovic, J.L. Scaffolding functions of arrestin-2 revealed by crystal structure and mutagenesis. Biochemistry 41, 3321–3328 (2002).

84. Lee, M.H., et al. The conformational signature of beta-arrestin2 predicts its trafficking and signalling functions. Nature 531, 665–668 (2016).

85. Nuber, S., et al. beta-Arrestin biosensors reveal a rapid, receptor-dependent activation/deactivation cycle. Nature 531, 661–664 (2016).

