## Supplementary information for "Small-molecule modulation of β-arrestins"

**Table of Contents:**


| | Page |
|---|---|
| S1: Methods | 3 |
| S2: Extended Data Figures and Tables | 17 |
| S3: References for Methods Section | 32 |

### Methods

#### Cell culture, antibodies, and reagents

HEK-293 cells (ATCC), including transient and stable lines, as well as CRISPR/Cas9 βarr1/2 knockout (KO) and parental lines[1], were maintained in Eagle's Minimum Essential Medium (MEM) supplemented with 10% fetal bovine serum (FBS) and 1% penicillin-streptomycin at 37°C and 5% $CO_2$. U2OS cells (DiscoveRx PathHunter) were cultured in MEM containing 2 mM L-glutamine, 10% FBS, and 1% penicillin-streptomycin under similar conditions. U2OS-based βarr recruitment and internalization assays were performed per the manufacturer's protocol (DiscoveRx, Fremont, CA). For chemokine-induced migration assays, leukocytes were isolated from wild-type mice as described previously[1-3]. Escherichia coli strains DH5α and BL21 (DE3) (New England Biolabs) were cultured in LB or Terrific Broth (Fisher Scientific) at 37°C. DH5α was used for plasmid amplification, and BL21 was used for recombinant protein expression. Sf9 insect cells (Spodoptera frugiperda, Expression Systems, Cat. No. 94-001F) were cultured in ESF 921 medium (Expression Systems, Davis, CA) at 27°C. Baculoviruses were generated and amplified according to the manufacturer's instructions. For serum starvation experiments, cells were cultured in serum-free medium supplemented with 0.1% BSA, 10 mM HEPES, and 1% penicillin-streptomycin. Transfections were performed using FuGene 6 (Promega) or Lipofectamine 3000 (Invitrogen), according to manufacturers' protocols. Monoclonal anti-FLAG M2-HRP (A8592) and anti-ERK1/2 (ABS44) antibodies were obtained from Sigma-EMD Millipore. HRP-conjugated secondary antibodies (NA9340-1ML and NA9310-1ML) were purchased from Cytiva. Protease and phosphatase inhibitor tablets (cOmplete™, PhosSTOP™) were obtained from Roche. Anti–phospho-p44/42 MAPK (Thr202/Tyr204) antibody (9101L) was sourced from Cell Signaling Technology.

#### Recombinant protein expression and purification

The expression and purification of βarrs have been described previously[4-7]. Briefly, *E. coli* BL21 (DE3) pLysS cells (New England Biolabs: C2527I) carrying pGEX4T1- *Rattus norvegicus* βarr1 or 2 construct, with their C-terminal truncated (at amino acid 393 or at 394, respectively), were cultured in Terrific Broth (Teknova, Hollister, CA) medium at

37°C. After $OD_{600}$ reached 0.6–0.8, the cells were induced with 0.1 mM isopropyl-β-D-thiogalactopyranoside (IPTG) at 18°C overnight. Bacteria were harvested by centrifugation (4000 RPM), and cell pellets were resuspended in lysis buffer [20 mM HEPES, pH 8, 150 mM NaCl, 10% glycerol, 1 mM EDTA, 0.2 mM dithiothreitol (DTT), 1 mM phenylmethylsulfonyl fluoride, and 1 mM benzamidine] at 4°C for 1 hr. Cell suspensions were sonicated, centrifuged (14,000 RPM, 30 min, 4°C), and the supernatant was loaded onto pre-equilibrated Glutathione-agarose resin (GoldBio). After 3 hours of binding at 4°C, the beads were washed, and thrombin was added for overnight cleavage. The proteins were further purified by anion exchange chromatography followed by size exclusion chromatography (SEC) using a Superdex 200 Increase 10/300 GL column on an ÄKTA FPLC system (GE Healthcare). Eluted fractions were analyzed by SDS–PAGE, pooled, concentrated using 30 kDa MWCO Amicon Ultra-15 Centrifugal Filter devices, flash-frozen in liquid nitrogen, and stored in aliquots at −80°C. The same rat βarr1 constructs with mutations (L129A, L129S, L129G, P131A, P133A, E134A, D135A, K138A, C140A, Y249A, C251A, E283A, K284A, R285A) were generated and purified in a similar manner (Extended Data Table 3). The heterotrimeric Gs protein complex, composed of Gαs, Gβ1, and Gγ2 subunits, was expressed in Spodoptera frugiperda (Sf9) cells using the baculovirus expression system and purified as previously described[8]. ERK2, c-Src, and p38α kinases were expressed and purified using previously established protocols[9,10]. Protein concentrations for each protein were determined by ultraviolet absorption at 280 nm and extinction coefficients estimated using the ExPASy ProtParam tool[11].

**Expression of $\beta_2$AR constructs in Sf9 cells using the baculoviral system**

The N-terminal FLAG-tagged T4 lysozyme fusion-$\beta_2V_2R$ construct ($\beta_2$AR residues 1–341 fused to $V_2R$ residues 328–372), bearing a TEV cleavage site, was co-expressed with GRK2-CAAX (membrane-anchored GRK2) or wild-type $\beta_2$AR in Sf9 insect cells using the Baculovirus Expression System, as described previously[4-6,12,13]. At 66 h post-infection, cells expressing T4L–$\beta_2V_2R$ were stimulated with 20 μM isoproterenol for 20 min at 37 °C to induce receptor phosphorylation (p$\beta_2V_2R$), while $\beta_2$AR-expressing cells remained untreated. All cells were washed extensively to remove residual agonist. For membrane preparation from these cells, briefly, cells were resuspended in cold homogenization buffer (75 mM Tris-HCl, pH 7.4, 2 mM EDTA, and cOmplete™ protease inhibitor) and collected by centrifugation at 500 × g for 5 minutes at 4°C. After two additional rounds of centrifugation at 500 × g for 5 minutes at 4°C, the supernatant was centrifuged at 21,000 × g for 30 minutes at 4°C to collect crude membrane fractions. Pellets were then washed with resuspension buffer (75 mM Tris-HCl, pH 7.4, 2 mM EDTA, 12.5 mM $MgCl_2$, cOmplete™ protease inhibitor, and PhosSTOP™ phosphatase inhibitor), passed through a 0.4 mm gauge needle 30 times using a syringe on ice, aliquoted, flash-frozen in liquid nitrogen, and stored at −80°C.

**Small molecule library and reagents**

A collection of structurally diverse, drug-like small-molecule libraries used in this work was obtained from the NCI/DTP Open Chemical Repository. The compound library comprised approximately 3.5K compounds, representing the structural diversity of over 250K unique small molecules. This DTP library included the NCI Diversity Set, natural products, and FDA-approved oncogenic drugs. Most compounds were certified as >95% pure by the supplier (NCI DTP Discovery Services). Powdered compounds were dissolved in 100% dimethyl sulfoxide (DMSO) and stored at −20°C. Isoproterenol (Sigma, I2760-1G), ICI-118,551, Carvedilol, angiotensin II (AngII; Sigma, A9525), and human epidermal growth factor (EGF) were purchased from Sigma-Aldrich, and [Arg8]-Vasopressin (AVP) was obtained from GenScript. All small molecule compounds, including BI-167107[14], were dissolved in DMSO and stored at −20°C as 100 mM stock solutions. The C-terminal peptide of the GPCR vasopressin-2 receptor ($V_2R$), known as $V_2Rpp$, was synthesized by the Tufts University Analytical Core Facility. Cellular agonist stimulations were performed at 37°C, as described in the figure legends.

**Differential scanning fluorimetry (DSF)**

A high-throughput differential scanning fluorimetry (DSF) assay was performed to identify βarr-binding small molecules from the drug-like compound library (DDLC) described above. The screen was conducted using the StepOnePlus™ Real-Time PCR System (Applied Biosystems) with the fluorescent reporter probe SYPRO Orange (Thermo Fisher

Scientific) in a 96-well format. Proteins were buffered in 20 mM HEPES (pH 7.5) with 100 mM NaCl. Small molecules (100 μM) were screened with either βarr1 or βarr2 (5 μM). DMSO was used as a vehicle control, while $V_2$Rpp (50 μM) served as a positive control. For DSF experiments involving the three positive control ligands ($V_2$Rpp, $IP_6$, and Heparin, each at 50 μM) binding to βarrs, HEPES buffer was used as the vehicle control. Excitation and emission filters for SYPRO Orange were set to 475 nm and 580 nm, respectively. The temperature was increased by 0.5°C every 30 seconds, from 25°C to 99°C, with fluorescence readings taken at each interval. Raw DSF data were analyzed using Applied Biosystems® Protein Thermal Shift™ Software. Fluorescence intensities were plotted as a function of temperature, and the midpoint of transition, or melting temperature ($T_m$), was calculated by plotting the first derivative of fluorescence emission as a function of temperature (dF/dT). The difference between the Tm of the protein-ligand complex and that of the protein alone represents the thermal shift ($\Delta T_m$), which indicates ligand binding to the protein of interest. For statistical analysis, experiments were conducted with at least three independent replicates per condition. The final selection of compounds for this study was guided by their efficacy in inhibiting βarr1/2 activity, along with favorable chemical properties, such as aqueous solubility and permeability, as described in the Results section. Three compounds were selected as candidate inhibitors: Cmpd-5 (NSC250682), (1S,4aR,5S,6S,6aR,9S,11aS,11bS,14R)-1,5,6,14-tetrahydroxy-4,4-dimethyl-8-methylenedecahydro-1H-6,11b-(epoxymethano)-6a,9-methanocyclohepta[a]naphthalen-7(8H)-one; Cmpd-46 (NSC302979), (Z)-3-ethoxy-6-hydroxy-4,4,13a-trimethyl-9-methylene-1,2,3,4,4a,5,6,9,10,11,12,13a-dodecahydro-7,10-(metheno)benzo[11]annulene-8,13-dione; and Cmpd-64 (NSC22070), (Z)-4,10a-dimethyl-7-methylene-8-oxo-1a,2,3,6,6a,7,8,9a,10,10a decahydrooxireno[2',3':8,9]cyclodeca[1,2-b]furan-6-yl acetate.

**Measurement of β-arrestin recruitment using the PathHunter assay**

βarr recruitment to the agonist-activated receptor ($\beta_2V_2R$) was measured using the DiscoveRx PathHunter β-arrestin assay[15], which employs enzyme fragment complementation. In this assay, the $\beta_2V_2R$ is fused to an inactive portion of β-galactosidase (ProLink™ tag), and βarr2 is fused to the complementary Enzyme Acceptor

(EA) portion, each stably expressed in U2OS cells. Upon agonist-induced recruitment of βarr2 to $\beta_2V_2R$, the β-gal fragments complement to form a functional enzyme, generating a measurable chemiluminescent signal. The signal intensity correlates directly with the extent of βarr2 recruitment. U2OS cells co-expressing $\beta_2V_2R$ and βarr2 were plated at a density of 25,000 cells per well in white, clear-bottom 96-well plates 24 hours before treatment. On the day of the experiment, cells were treated with either a single dose (50 μM) or varying concentrations of a specific βarr modulator or vehicle control in Hanks' balanced salt solution (Sigma-Aldrich) with 20 mM HEPES (pH 7.4) and 0.05% BSA. Cells were incubated at 37°C, 5% $CO_2$, and ~100% relative humidity for ~30 minutes, followed by stimulation with either 10 nM isoproterenol (ISO) or a serial dilution of ISO for 60 minutes at 37°C. After agonist stimulation, PathHunter reagents were added, and cells were incubated for another 60 minutes at ambient temperature. Luminescence signals were measured using a CLARIOstar microplate reader (BMG Labtech).

**Measurement of β-arrestin recruitment using the NanoBiT assay**

To elucidate the effects of compounds on the individual interactions of βarr1 and βarr2 with the receptor, NanoLuc Binary Technology (NanoBiT) assays were performed using HEK293 CRISPR/Cas9 βarr1/2 knockout (KO) cells transiently transfected with pcDNA 3.1 $V_2R$-LgBiT for the receptor and either pcDNA 3.1 βarr1-SmBiT or pcDNA 3.1 βarr2-SmBiT constructs. In this NanoBiT βarr recruitment assay, the SmBiT tag was placed on the N-terminus of human βarr1/2, and the LgBiT tag was fused to the N-terminal region of human $V_2R$, which also included an HA signal sequence and a FLAG tag (DYKDDDDK) on the N-terminus of $V_2R$, as previously described[16-18]. Cells were plated at a density of 100,000 cells per well in a white, clear-bottom 96-well plate one day before transfection. The transfection solution was prepared by combining $V_2R$-LgBiT and either βarr1-SmBiT or βarr2-SmBiT constructs in a 1:5 ratio (25 ng of receptor and 125 ng of βarr1/2). Transfections were performed using Lipofectamine 3000 (Invitrogen, catalog #L3000015) according to the manufacturer's instructions. Forty-eight hours post-transfection, cells were washed and incubated in Opti-MEM™ for 60 minutes at 37°C before conducting NanoBiT assays. Coelenterazine-h and individual modulators (compounds 5, 46, or 64) were added to cells at a final concentration of 50 μM, approximately 30 minutes prior to measurement. After establishing a baseline response for 2 minutes, cells were stimulated with 100 nM AVP or vehicle, and luminescence was measured for an additional 20 minutes. The complemented (SmBiT + LgBiT) NanoLuc signals were detected at 550 nm using a PHERAstar FSX instrument (BMG LabTech). Ligand-induced changes in luminescent units were normalized to baseline and vehicle, and the area under the curve (AUC) for the entire time course was calculated to determine the cumulative agonist-induced response and modulator effects. This approach allowed for a robust determination of the maximal effect of modulators on βarr1/2 recruitment to AVP-activated $V_2R$.

**FRET-based cAMP accumulation measurement**

To measure cellular cAMP production in live cells mediated by stimulatory G protein, Gαs-coupled $\beta_2AR$ activation, FRET-based Epac sensors were used as described previously[19]. The Epac2 (ICUE2) sensor contains a CFP and YFP FRET pair. HEK293 cells stably expressing ICUE2 were plated in poly-D-lysine-coated, black, clear-bottom 96-well plates (Corning) at a density of 50,000 cells per well. At least 16 hours after plating, cells were washed with PBS and incubated in HEPES-buffered saline solution (10 mM HEPES, 150 mM NaCl, 5 mM KCl, 1.5 mM $MgCl_2$, 1.5 mM $CaCl_2$, 10 mM glucose, 0.2% BSA, pH 7.4) for one hour at 37°C. Cells were then treated with either βarr small-molecule modulators (40 μM) or DMSO for 5 minutes, and baseline fluorescence was monitored. Real-time cAMP measurement was initiated by stimulating cells with 10 μM isoproterenol (ISO) at 37°C. FRET changes corresponding to cAMP accumulation were measured as changes in the background-subtracted 480 nm/535 nm fluorescence emission ratio (CFP/YFP), reflecting changes in cAMP levels. The entire cAMP accumulation profile was quantified by calculating the area under the curve (AUC) for the time course. To assess the effect of modulators on the agonist-induced cAMP response, AUC values were expressed as a percentage of the response to ISO in the presence of DMSO (set as 100%), enabling comparison of the effect of each modulator relative to this agonist-alone control.

**Intracellular calcium measurement**

Intracellular [$Ca^{2+}$] release was measured using the FLIPR Calcium 6 assay kit with a FlexStation 3 microplate reader, following the manufacturer's instructions (Molecular Devices, LLC) and as described previously[20]. Briefly, HEK293 cells stably expressing human Angiotensin II type 1 receptor ($AT_1R$), parental HEK293 cells transiently expressing $AT_1R$, or HEK293 CRISPR/Cas9 βarr1/2 KO cells transiently expressing $AT_1R$ were seeded in poly-D-lysine-coated, black 96-well assay plates at a density of 40,000 cells per well and incubated for 24 hours. On the day of the experiment, cell plates were loaded with FLIPR Calcium 6 reagents and treated with βarr small-molecule modulators (10 μM) or vehicle (DMSO) for 30 minutes. After establishing basal fluorescence ($F_0$), the cells were treated with the agonist AngII (30 pM for stably expressing $AT_1R$ or 120 pM for transiently expressing $AT_1R$) while fluorescence (F) intensity was monitored in real time. The entire calcium transient profile was quantified by calculating the area under the curve (AUC) for the time course. To assess the effect of modulators on the agonist-induced $Ca^{2+}$ responses, AUC values were expressed as a percentage of the response to AngII in the presence of buffer alone (set as 100%), enabling comparison of each modulator's effect relative to this agonist-alone control.

**Measurement of receptor internalization by PathHunter assay**

βarr-mediated receptor internalization was measured using the DiscoveRx PathHunter active receptor endocytosis assay according to the manufacturer's protocol (DiscoveRx, Fremont, CA) and as previously described[15]. Briefly, $\beta_2V_2R$ was transiently transfected into U2OS cells stably expressing an Enzyme Acceptor (EA)-tagged βarr2 and an endosome-localized ProLink-tagged protein. The next day, cells were seeded at 25,000 cells per well in white, clear-bottom 96-well assay plates and incubated for 24 hours before the experiment. On the day of the experiment, cells were treated with varying concentrations of a specific βarr modulator or vehicle control in Hanks' balanced salt solution (Sigma-Aldrich) with 20 mM HEPES (pH 7.4) and 0.05% BSA. Cells were incubated at 37°C, 5% $CO_2$, and ~100% relative humidity for ~30 minutes, followed by stimulation with a series of concentrations of agonist (ISO) for 60 minutes at 37°C. After agonist stimulation, PathHunter reagents were added, and cells were incubated for another 60 minutes at ambient temperature. Receptor–βarr complex internalization was detected as luminescence resulting from the complementation of β-gal fragments (Enzyme Acceptor and ProLink) within endosomes. Luminescence signals were measured using a CLARIOstar microplate reader (BMG Labtech).

**BRET based receptor internalization assay**

BRET-based assays were conducted to measure receptor internalization using a bystander BRET format as previously described[21]. Briefly, HEK293 cells transiently expressing $V_2$R-RLucII (BRET donor) and the early endosome marker 2xFYVE-mVenus (BRET acceptor) were pretreated with vehicle or βarr modulator (40 μM) for 30 minutes. Receptor association with the endosome marker was measured as a BRET signal after stimulation with a range of AVP concentrations. BRET measurements were performed using the Synergy2 (BioTek®) microplate reader with filter sets of 410/80 nm and 515/30 nm to detect RLucII (donor) and mVenus (acceptor) emissions, respectively. The BRET signal was calculated as the ratio of light intensity emitted by the acceptor over the donor, and the 'net BRET' ratio was determined by subtracting the vehicle control ratio from the corresponding AVP-treated ratio.

**Chemotaxis assays**

Chemotaxis assays were performed similarly to previously described protocols [2]. Briefly, murine T cells were isolated from the spleens of wild-type mice, subjected to erythrocyte lysis, and filtered through a 70-μm filter. The cells were then suspended in RPMI 1640 medium containing 0.5% BSA and treated for 30 minutes with βarr modulators or vehicle. A total of $1 \times 10^6$ cells in 100 μL of medium were added to the upper chamber of 6.5-mm diameter, 5-μm pore polycarbonate Transwell filters (Corning Costar), and cells migrated toward 100 nM CCL19 in the lower chamber for 2 hours at 37°C. T cells that migrated to the lower chamber were collected, resuspended, washed, and stained for flow cytometry analysis using a Live/Dead marker (Aqua Dead, ThermoFisher) and antibodies for cell surface markers ($CD45^+$, $CD3^+$, $CD4^+$, and $CD8^+$) prior to paraformaldehyde fixation. The total live T cell population (CD45+ and CD3+) and subset populations ($CD45^+$, $CD3^+$, and $CD4^+$, or $CD45^+$, $CD3^+$, and $CD8^+$) were measured using a BD LSR Fortessa machine from the Flow Cytometry Shared Resource (FCSR) at the Duke Cancer Institute (Durham, NC). CountBright beads (ThermoFisher) were added immediately after resuspension of the lower chamber contents to correct for volume differences and any cell loss during wash steps. Percent migration was calculated by determining the percentage of migrated cells treated with compounds relative to control wells, with migration in CCL19-treated wells alone set as 100%. The use of mouse splenocytes *ex vivo* for this T cell migration was conducted under institutional guidelines for the care and use of laboratory animals. No live animal procedures were performed. A representative gating tree is shown in Extended Data Fig. 5.

**Isothermal titration calorimetry**

Isothermal Titration Calorimetry (ITC) measurements were performed on a MicroCal iTC200 system (Malvern, PA) at 25°C. Prior to ITC experiments, all proteins (βarr1, βarr2, or βarr1 mutants) were extensively dialyzed against 20 mM HEPES (pH 7.4), 100 mM NaCl, and 3 μM TCEP. Protein concentrations were determined by spectrophotometry at 280 nm, using extinction coefficients calculated from each protein sequence via the ProtParam program[11]. The dialysis buffer was used to dilute DMSO stock solutions of βarr small molecule modulators to their final concentrations for measurements. For each ITC experiment, βarr modulators (Cmpd-5 at 350 μM, Cmpd-46 at 450 μM, and Cmpd-64 at 450 μM) were loaded into the syringe and titrated into the calorimetric cell containing βarr1, βarr1 mutants, or βarr2 (at approximately 30 μM, 20 μM, 40 μM, and 40 μM respectively, for each modulator). The reference cell was filled with distilled water. In all experiments, the titration sequence typically consisted of an initial 0.4 μl injection, followed by 19 injections of the βarr modulator compound at 2 μl each, with a 180-second interval between injections to allow thermal power to return to baseline. During the experiment, the reference power was set to 7 $\mu cal \cdot s^{-1}$, and the sample cell was stirred continuously at 750 rpm. Raw data, excluding the peak from the first injection, were baseline-corrected, integrated, and normalized. Data were analyzed and fit using a one-site independent-binding model to obtain the equilibrium dissociation constant, $K_D$ (from the association constant $K_A = 1/K_D$), stoichiometry (N), and thermodynamic parameters such as enthalpy (ΔH) and entropy (-TΔS) of binding. Data were analyzed using MicroCal PEAQ-ITC analysis software (version 1.1.0.1262, Malvern, PA).

**Radioligand binding experiments**

To assess the effects of βarr modulators on βarr- or Gαs–βγ-promoted high-affinity agonist binding, [$^3$H]-methoxyfenoterol ([$^3$H]-Fen) binding assays were performed using membranes from Sf9 cells expressing phosphorylated $\beta_2V_2R$ (p$\beta_2V_2R$) or $\beta_2AR$ (Ahn et al., 2017). Membranes were prepared from Sf9 cells co-expressing FLAG-tagged T4L–$\beta_2V_2R$ and GRK2-CAAX under agonist stimulation, or unstimulated $\beta_2AR$ alone as described above[15]. Reactions (150 µL) contained 6 nM [$^3$H]-Fen (12.6 Ci/mmol), p$\beta_2V_2R$ membranes, βarr1/2 or βarr1 mutants (2 µM), and βarr modulators (100 µM) or vehicle in HN100 buffer (20 mM HEPES, pH 7.4, 100 mM NaCl, 12.5 mM $MgCl_2$). For Gαs–βγ assays, $\beta_2AR$ membranes were incubated with 4.3 nM [$^3$H]-Fen and Gαs–βγ heterotrimer (200 nM) in G protein assay buffer with or without modulators. Nonspecific binding was defined using propranolol (25 µM). Following 90 min incubation at room temperature, reactions were filtered onto PEI-soaked GF/B filters and washed with cold buffer. Bound [$^3$H]-Fen was extracted overnight in scintillation fluid and quantified by scintillation counting. Specific binding was calculated by subtracting nonspecific from total binding.

**Cytotoxicity assay**

MTT assay was performed according to the manufacturer's instructions (Roche Diagnostics GmbH, Germany). HEK293 and U2OS cells were seeded in 96-well plates. The following day, cells were treated with the indicated concentration of modulator or vehicle for 8 h. To evaluate the cytotoxic effects of the compounds, cells were incubated with 3-(4,5-dimethylthiazol-2-yl)-2,5-diphenyltetrazolium bromide (MTT) reagent at 37 °C for 4 h. The optical density (OD) was measured at 595 nm (the absorbance of each sample was measured at 560 and 670 nm). The optical density values of blue formazan formed in live cells based on the reduction of MTT were determined at 595 nm. Cell viability was expressed as the percentage of MTT reduction in compound-treated cells compared to vehicle-treated cells.

**Live-cell ERK1/2 activation assay**

To evaluate the effect of small-molecule modulators on βarr-dependent ERK1/2 signaling, HEK293 cells stably expressing $\beta_2$AR were serum-starved for 6 h[22,23], then pretreated with βarr modulators (30 µM) or vehicle for 20 min at 37 °C. Cells were subsequently stimulated with carvedilol (10 µM, 5 min), a βarr-biased agonist known to promote ERK1/2 phosphorylation via βarr scaffolding of RAF–MEK–ERK signaling components. Cells were lysed, sonicated (15 sec), and centrifuged at 14,000 × g for 15 min at 4 °C. Equal amounts of total protein were resolved by SDS-PAGE on 4–20% Tris-Glycine gels (Thermo Fisher Scientific, Carlsbad, CA), transferred to nitrocellulose or PVDF membranes, and immunoblotted with anti–phospho-ERK1/2 (1:2,000; Cell Signaling) and total ERK1/2 (1:10,000; Millipore-Sigma). HRP-conjugated secondary antibodies were used for detection. Protein bands were visualized using Pierce SuperSignal West Pico ECL substrate (Thermo Fisher Scientific, Waltham, MA), imaged on a ChemiDoc XRS system (Bio-Rad Laboratories, Hercules, CA), and quantified by densitometry using Image Lab (Bio-Rad). Data were analyzed using GraphPad Prism.

**Cryo-EM sample preparation and data acquisition**

The expression construct for βarr1-BRIL was designed by inserting the thermostabilized apocytochrome b562 (M7W, H102I, K106L) from *Escherichia coli* (BRIL) between residues 176-182 at the hinge region of rat βarr1[7] truncated at residue 393 and purified as wild-type βarr1. βarr1-BRIL was incubated with 1.5-fold molar excess of anti-BRIL Fab (BAG2)[24] and 2-fold molar excess of anti-Fab-Nb (aFabNb)[25] for 1 hour at room temperature. The complex was subjected to SEC on a Superdex 200 Increase column (Cytiva Life Sciences) in 20 mM HEPES 7.5, 150 mM NaCl buffer. Peak fractions were concentrated to 2 mg/ml using Vivaspin® 6 column with molecular weight cut-off of 30,000 kDa (Sartorius). The complex was incubated for 2 hours on ice with 50-fold molar excess of Cmpd-5 solubilized in DMSO (βarr1-BRIL–BAG2–aFabNB–Cmpd-5) or equivalent amount of DMSO (βarr1-BRIL–BAG2–aFabNB–Apo), and then concentrated to 7-8 mg/ml. The sample was applied to glow-discharged 300-mesh holey-carbon grids

(Quantifoil R1.2/1.3, Electron Microscopy Sciences) using a Vitrobot Mark IV (Thermo Fisher Scientific) at 4 °C and 100% humidity. The data were collected on a Titan Krios transmission electron microscope (Thermo Fisher) operating at 300 kV equipped with a K3 direct electron detector (Gatan) in counting mode with a BioQuantum GIF energy filter (slit width of 20 eV) at a magnification of ×81,000 corresponding to a pixel size of 1.08 Å at the specimen level. 60-frame movies with a dose rate of ~15 electrons per pixel per second and a total accumulated dose of ~54-60 electrons per $Å^2$ were collected using the *Latitude-S* (Gatan) single-particle data acquisition program. The nominal defocus values were set from −0.8 to −2.5 μm.

**Cryo-EM data processing**

Movies were subjected to beam-induced motion correction using Patch Motion Correction in CryoSPARC v4.0.1 [26] followed by determination of Contrast transfer function (CTF) parameters in Patch CTF. Micrographs with CTF fit better than 3.5 Å were used for further analysis. Particles manually selected from 15 micrographs were used to train a model in the particle picking tool Topaz v0.2.5a [27]. The trained model was used to pick particles in all micrographs generating 1,572,529 particle projections for βarr1-BRIL–BAG2–aFabNB–Cmpd-5 and 1,307,318 particle projections for βarr1-BRIL–BAG2–aFabNB–Apo. The particles were rescaled to the pixel size of 1.3824 Å for further processing. A subset of particles (500,000) was subjected to Ab Initio model generation with 5 classes. All particles were then subjected to five rounds of heterogeneous refinement. The resulting particle stacks (396,911 particles for βarr1-BRIL–BAG2–aFabNB–Cmpd-5 and 832,520 particles for βarr1-BRIL–BAG2–aFabNB–Apo) were used in non-uniform refinement and local refinement with a mask excluding aFabNb and the constant region of BAG2. Then particles were subjected to 3D classification without alignment in CryoSPARC with a mask excluding aFabNb and the constant region of BAG2. Finally, the best classes with 177,430 particles (βarr1-BRIL–BAG2–aFabNB–Cmpd-5) and 479,224 particles (βarr1-BRIL–BAG2–aFabNB–Apo) were subjected to another round of local refinement in CryoSPARC, generating a map with a global resolution of 3.47 Å and 3.52 Å, respectively. The Cryo-EM maps were post-processed using DeepEMhancer with highRes deep learning model[28].

**Model building and refinement**

The initial models were built manually by fitting the crystal structure of βarr1 (PDB: 1G4M)[29] into the experimental electron densities using UCSF Chimera v1.15[30]. The BRIL-BAG2-aFabNb part of the model was derived from the Cryo-EM structure of Frizzled5-BRIL-BAG2-aFabNb (PDB: 6WW2)[31]. The structures were refined by combining manual adjustments in Coot v0.9.8.3[32] and ISOLDE[33] in UCSF ChimeraX 1.6.1[30], followed by real-space refinement in PHENIX v1.20.1-4487[34] with Ramachandran, rotamer, torsion, and secondary structure restraints enforced. The models were validated with MolProbity v4.5.1[35]. Cryo-EM refinement statistics are summarized in Extended Data Table 1.

**Modeling and molecular dynamics (MD) simulations**

The apo βarr1 model in the basal state and the $\beta_2V_2R$–βarr1 complex (PDB code: 6TKO)[36] model were built and simulated as described previously[37]. Due to a significant number of missing residues in the Cmpd5-bound βarr1 cryo-EM structure, we constructed a Cmpd5-bound βarr1 model by placing Cmpd5 in the MCL cleft of an equilibrated basal βarr1 model according to the experimentally derived location of the bound Cmpd5. A βarr1 model in the active state was constructed by homology modeling with Modeller (v10.0)[38] using the crystal structure of the $V_2$Rpp-bound βarr1 in the active state (PDB code: 4JQI)[5] as the template. Parts without any template, i.e., residues 1 to 5 and 307 to 311 were *ab initio* modeled with Modeller. The resulting model with the lowest DOPE score was selected for the following steps. The selected βarr1 models were processed by the Protein Preparation Wizard of Schrodinger Suite (version 2023-1). The first and last residues of this model are in positively and negatively charged states, respectively, without capping, as assumed in their natural condition. The prepared model was then immersed in a simulation water box using the System Builder of Schrodinger Suite (version 2023-1). A simple point charge (SPC) water model was used to solvate the system, and $Na^{+}$, and $Cl^{-}$ ions were added to neutralize the system and the salt concentration of the system was increased to 0.15 M. The total system size was ~148,000 atoms.

MD simulations were carried out using Desmond MD System (version 6.1; D.E. Shaw Research, New York, NY)[39] with the OPLS4 force field[40]. The simulation systems were minimized and equilibrated with restraints on the ligand heavy atoms and the protein backbone. For both the equilibrations and the following production runs, the constant temperature 310 K was maintained by Langevin dynamics, 1 atm constant pressure was achieved with the Langevin piston method[41]. A cutoff distance of 9 Å was used for the nonbonded interactions, and the particle-mesh Ewald summation method was used for the electrostatics interactions. The integration timestep was set to 2.5 fs. The isothermal-isobaric (NPT) ensemble was used in a periodic boundary condition. The production runs of βarr simulations are without any restraints. The analysis and visualization were performed with VMD[42] and PyMol (Schrodinger).

**Graphing and statistical analyses**

All graphs were generated and analyzed using GraphPad Prism 10.0 (GraphPad Software, Inc., La Jolla, CA). Dose-response curves were fitted to a log(agonist) versus response model with parameters for span, baseline, and $EC_{50}$, and minimum baseline was corrected to zero. For statistical comparisons, one-way ANOVA with Dunnett's post hoc test was generally used for comparisons across more than two groups, and two-way ANOVA with Sidak’s post hoc test or similar was used for comparisons involving multiple conditions, as specified in the figure legends. Most experiments were conducted with three biological replicates, and additional replicates served as controls. Replicates in the figure legends refer to biological replicates, with technical replicates included in some experiments for intra-replicate variation. Differences with $P$-values $< 0.05$ were considered significant. Further statistical details and replicate information are provided in the figure legends.

### Extended Data Figures and Tables

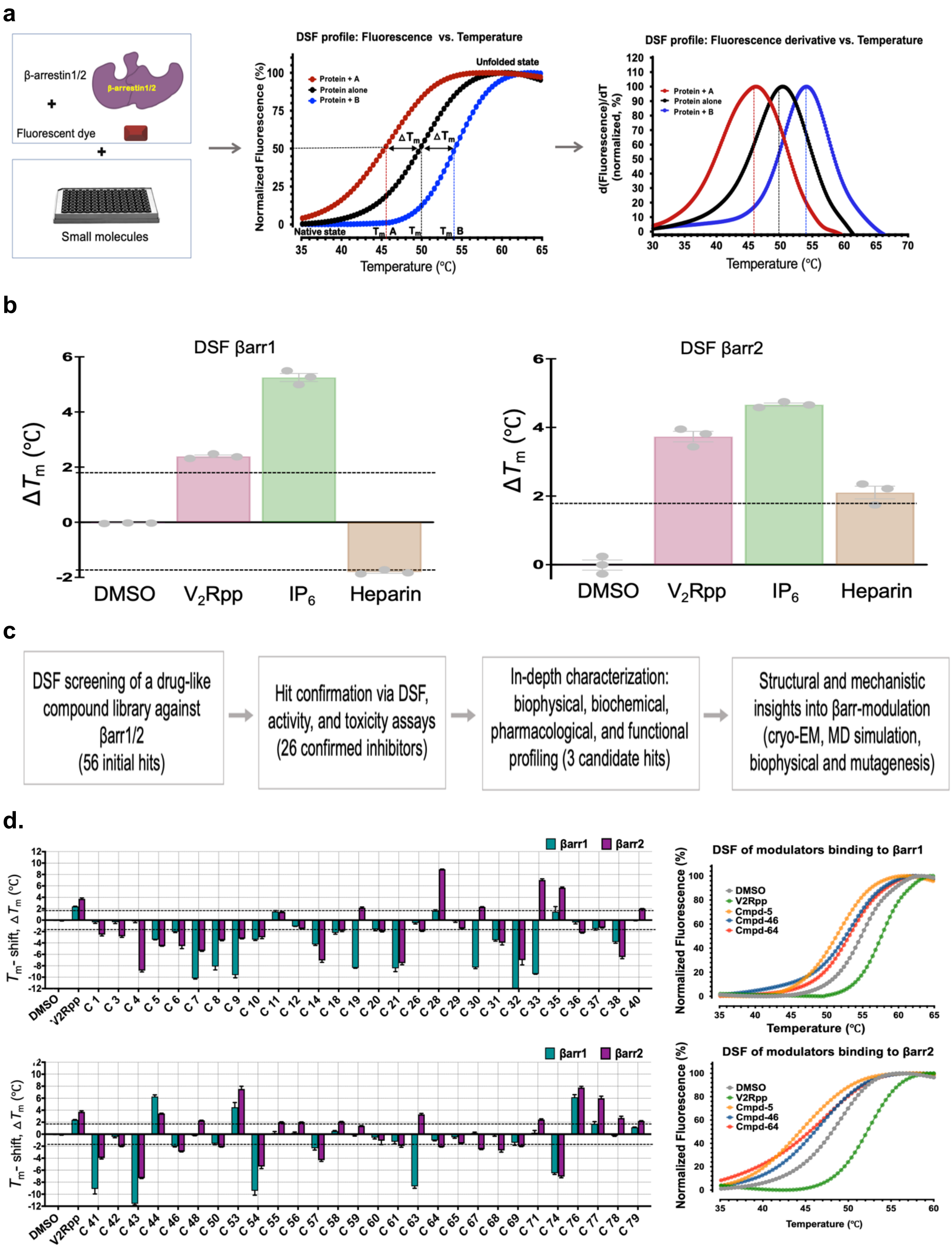

**Extended Data Fig. 1 |** Differential scanning fluorimetry (DSF) platform for screening β-arrestin1/2 binding small molecules. **a**, Schematic of the DSF workflow for screening βarr1/2 binders. DSF of βarrs (5 μM) was performed using a fluorescent dye, with fluorescence measured as a function of temperature. Representative DSF profiles are shown as normalized fluorescence (middle) and first derivative fluorescence (dF/dT) (right) as a function of temperature, indicating melting transitions ($T_m$) and ligand-induced thermal shifts ($\Delta T_m$). **b,** DSF assay validation using known βarr ligands–$V_2$Rpp, $IP_6$, and heparin (50 μM each)–showing $\Delta T_m$ for βarr1 (left, ~54.5 °C) and βarr2 (right, ~49 °C). $T_m$ shifts ($|\Delta T_m| \geq 1.8$ °C, dashed lines) denote significant binding, with the threshold informed by the $\Delta T_m$ of βarr1 upon heparin binding. Data are mean ± s.e.m., n = 3. **c**, Screening workflow outlining the multi-tiered process: initial DSF screening of a compound library (56 preliminary hits), followed by hit confirmation (26 validated binders), in-depth characterization yielding three candidate βarr modulators, and mechanistic and structural studies to elucidate modes of action. **d**, DSF data for βarr1 and βarr2 (5 μM) in the presence of screened small molecules (100 μM, n = 3). Bar plots show $\Delta T_m$ for each compound, highlighting potential binders ($\Delta T_m$; stabilizing and destabilizing). Small molecules with $|\Delta T_m| \geq 1.8$ °C were considered binders. Right, representative DSF melting curves depict binding profiles of select modulators compared to controls. $V_2$Rpp served as a positive control. Data are mean ± s.e.m., n = 3.

**a**

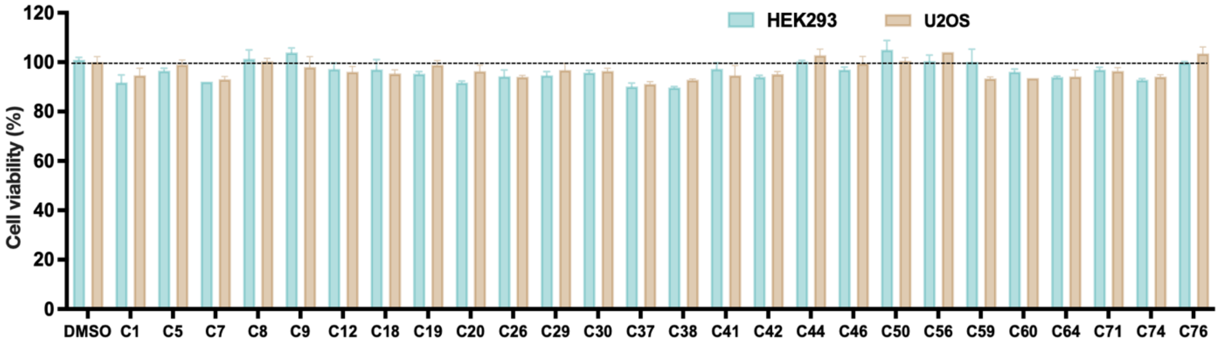


**b**

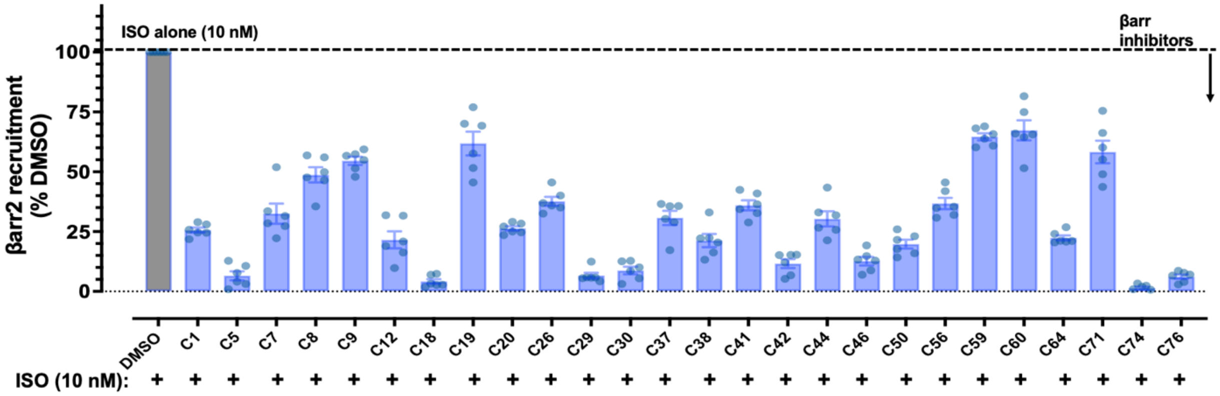

**c**

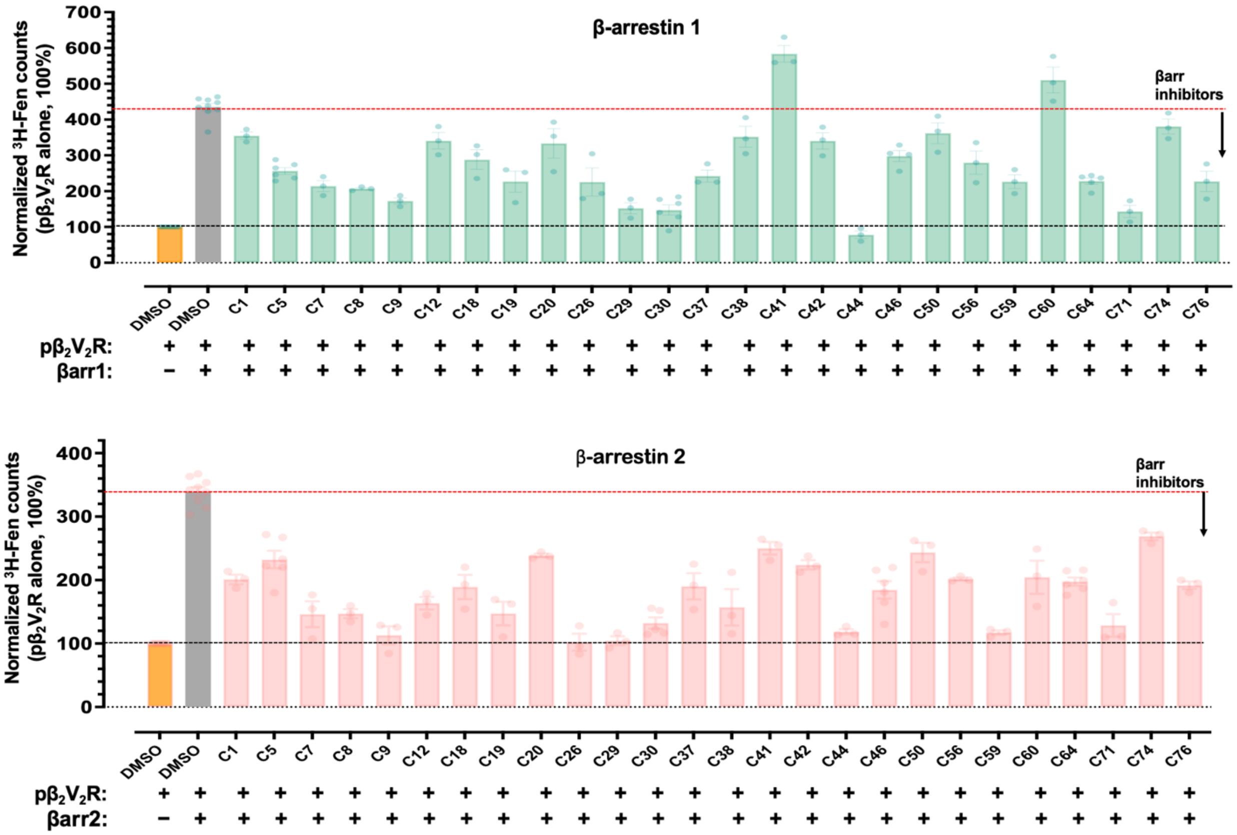


**Extended Data Fig. 2 |** Characterization of putative β-arrestin modulators in cell-based and in vitro assays. **a**, Cell viability of HEK293 and U2OS cells treated with 50 µM of each compound or vehicle (DMSO) for 8h, assessed using an MTT assay. Viability is expressed as a percentage relative to DMSO control (dashed line) (HEK293, light green; U2OS, beige). Data are mean ± s.e.m., n = 3. **b**, Effect of putative βarr modulators on βarr2 recruitment to agonist-activated $\beta_2V_2R$. DiscoveRx-U2OS cells expressing βarr2 and $\beta_2V_2R$ were treated with each compound (50 µM, 30 min) before stimulation with isoproterenol (10 nM). Recruitment was quantified as a percentage of DMSO control. Signals below the dashed line (agonist-only control) indicate inhibition of βarr2 recruitment. Data are mean ± s.e.m., n = 5. **c**, Effect of modulators on βarr-promoted high-affinity agonist binding to phosphorylated $\beta_2V_2R$ ($p\beta_2V_2R$). *In vitro* radioligand binding assays using Sf9 membranes with $p\beta_2V_2R$ assessed agonist ([$^3$H]-Fenoterol, 6 nM) binding in the presence of βarr1 (top) or βarr2 (bottom), with or without modulators (100 µM). The first bar (orange) shows $p\beta_2V_2R$ alone, while the second bar (grey) represents $p\beta_2V_2R$ with βarr (2 µM). Modulator-induced decreases in binding (below the dashed red line) reflect the inhibition of βarr-mediated receptor engagement. In summary, secondary DSF screens, toxicity assays, and functional βarr activity tests were used to refine initial hits, prioritizing compounds with favorable chemical and pharmacological properties. Data are mean ± s.e.m., n = 3.

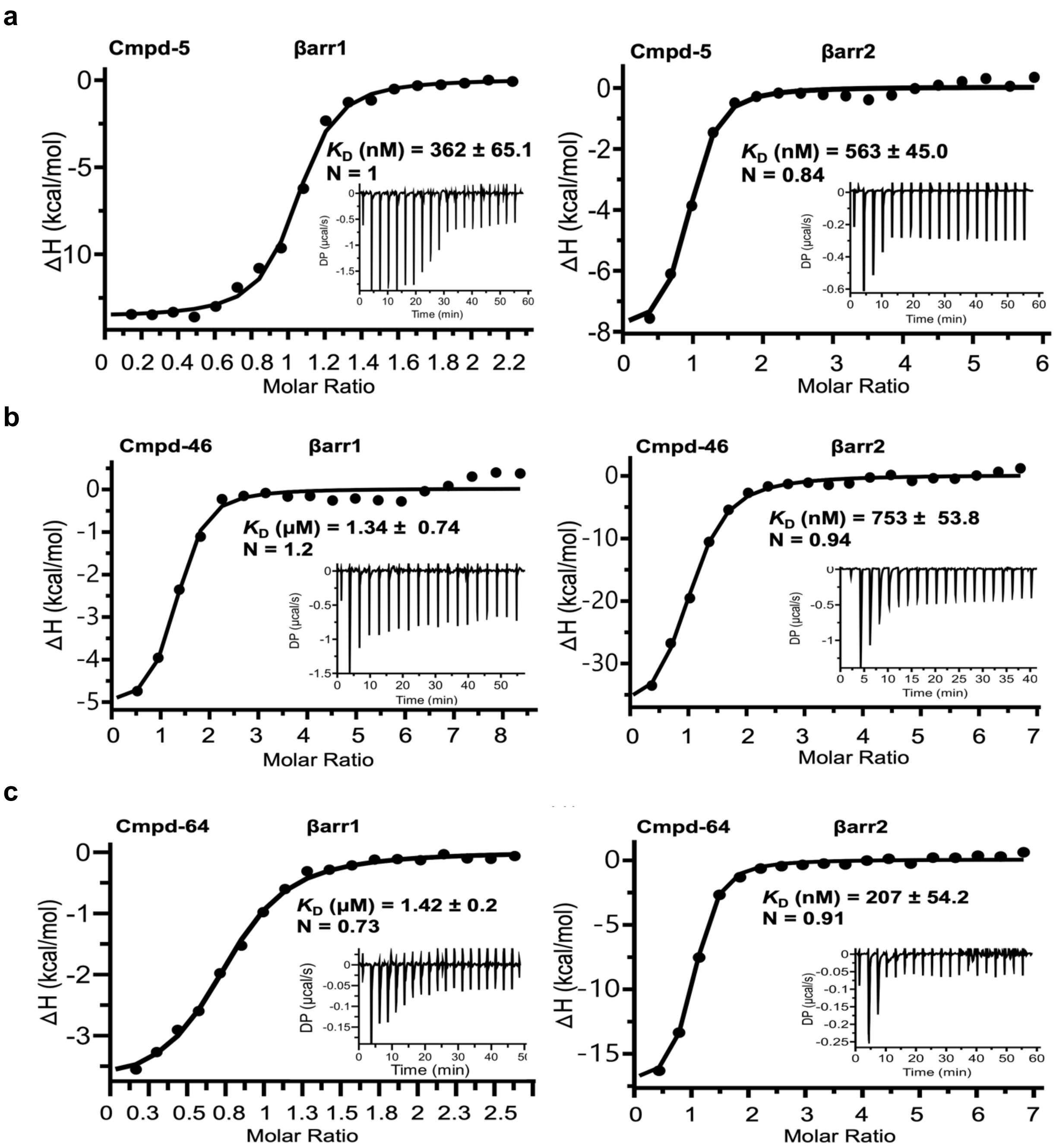


**Extended Data Fig. 3** | Isothermal titration calorimetry (ITC) analysis of modulator binding to βarr1 and βarr2. **a–c**, ITC isotherms for modulator binding to βarr1 (left) and βarr2 (right). **a**, Cmpd-5; **b,** Cmpd-46; **c,** Cmpd-64. Raw injection heats (insets) and fitted binding isotherms (solid lines) were modeled using a one-site independent binding model. Apparent Equilibrium dissociation constants ($K_D$) and stoichiometry values (N) are shown as mean ± s.e.m., based on fits from independent measurements. Representative titration curves from independent experiments are shown. Cmpd-5 bound both isoforms comparably; Cmpd-46 showed slight βarr2 preference, and Cmpd-64 exhibited ~6-fold selectivity for βarr2 over βarr1. Stoichiometry values for most interactions were consistent with 1:1 binding.

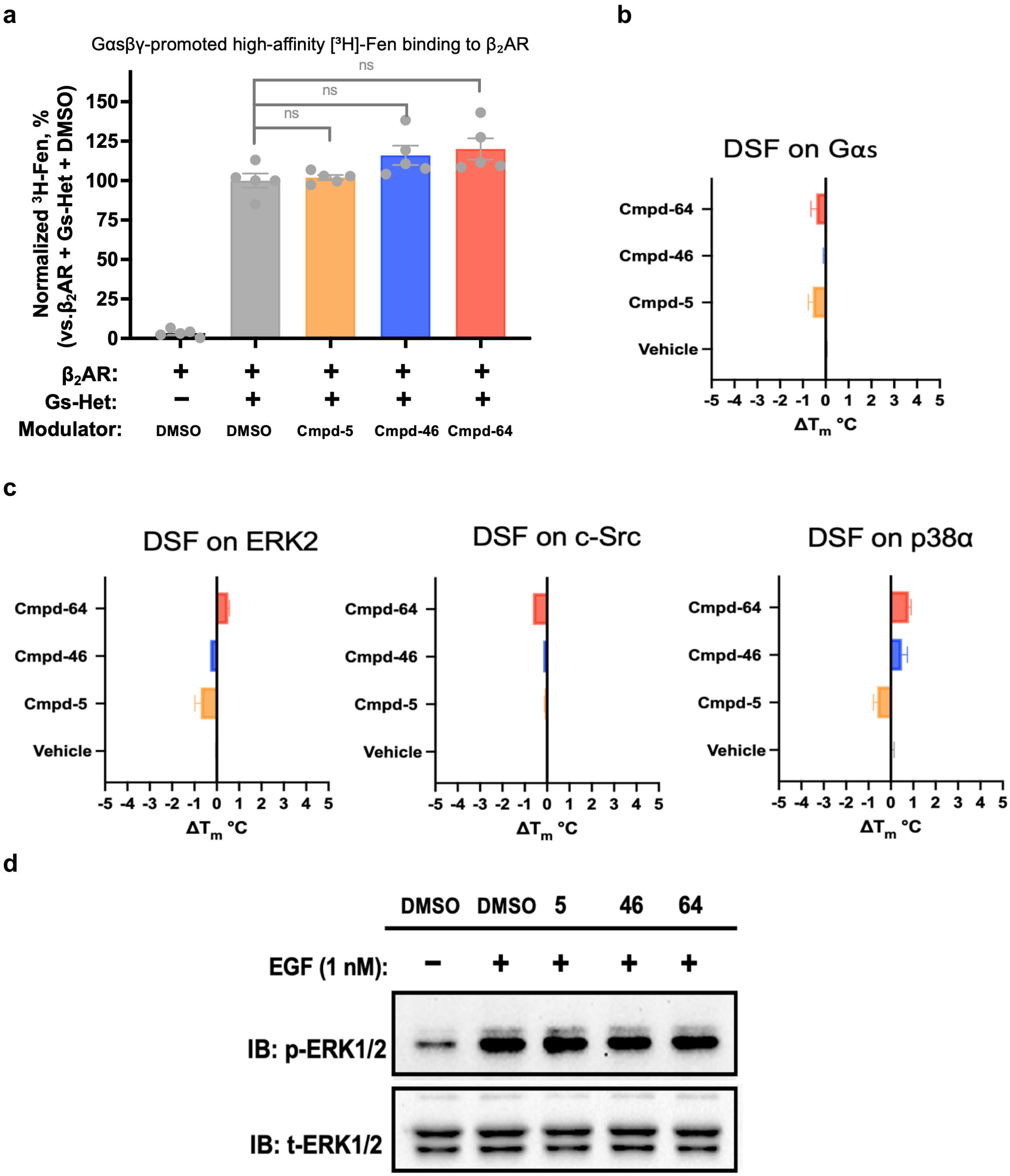


**Extended Data Fig. 4** | β-arrestin modulators display target specificity with minimal off-target effects. **a**, βarr modulators do not alter Gαs heterotrimer-mediated high-affinity agonist binding to β2AR. Radioligand binding assays were performed in Sf9 membranes expressing β2AR using [$^{3}$H]-fenoterol (4.3 nM) in the presence of purified Gαs–βγ (200 nM), with or without modulators (Cmpd-5, -46, or -64; 100 μM). Gαs–βγ increased agonist binding (grey), while modulator-treated samples showed no significant change. Data are normalized to $\beta_2$AR + Gαs–βγ + DMSO. Mean ± s.e.m., n = 5. **b**,**c**, Differential scanning fluorimetry (DSF) assays with purified GST–Gαs (5 µM) (b) and kinases ERK2, c-Src, and p38α (5 µM each) (c) revealed negligible shifts in melting temperature ($\Delta T_m$) upon modulator treatment, indicating minimal off-target engagement with Gαs or representative kinases. Mean ± s.e.m., n = 3. **d**, βarr modulators do not interfere with EGFR-mediated ERK activation. HEK293 cells were treated with modulators (30 µM) or DMSO, followed by EGF (1 nM). Immunoblots of phosphorylated and total ERK1/2 showed no modulation of ERK signaling in response to treatment.

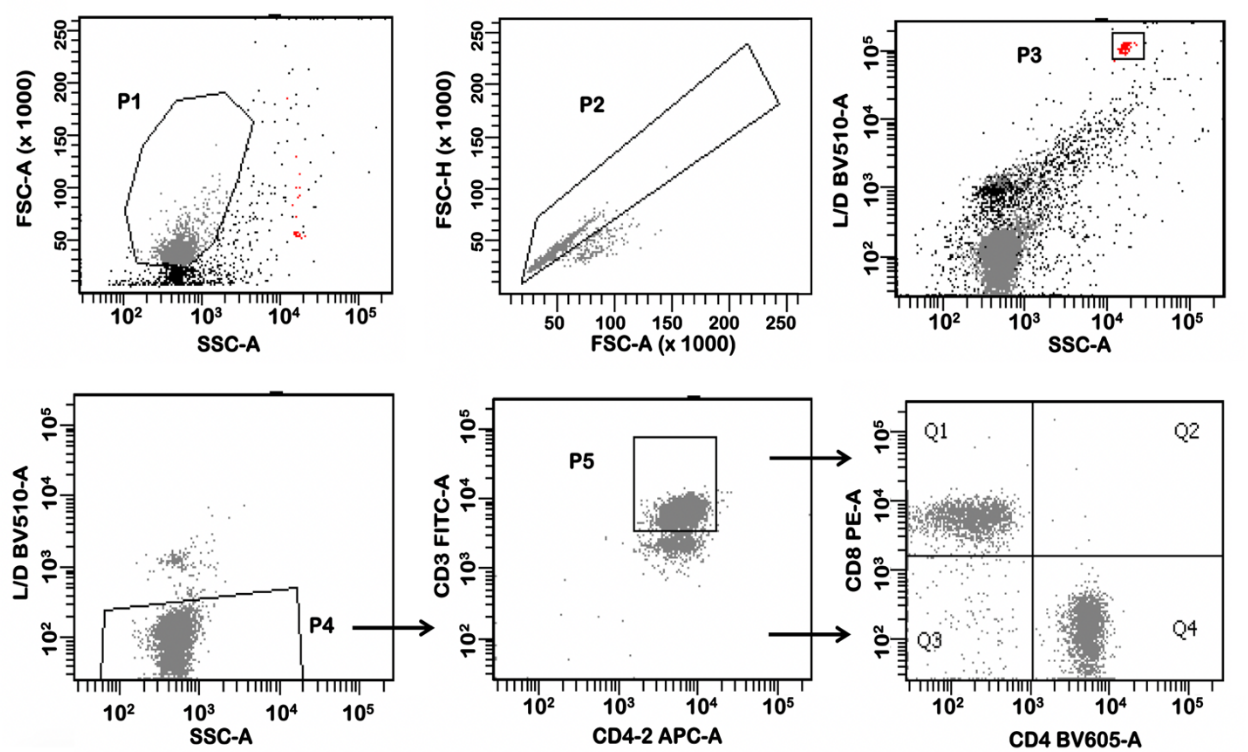


**Extended Data Fig. 5** | General gating strategy for T cell migration assay analysis by flow cytometry. Gating strategy for the analysis of the T cell migration assay. A representative plot is shown for DMSO-treated mouse splenocytes input into a transwell and allowed to migrate in response to CCL19 stimulation. **p1**, forward scatter area (FSC-A) and side scatter area (SSC-A); **p2**, forward scatter height (FSC-H) and forward scatter area (FSC-A); **p3**, counting beads; **p4**, live/dead aqua and side scatter area (SSC-A); **p5**, CD3 FITC and CD45.2 APC; **q1**, CD8 PE; q4, CD4 BV605.

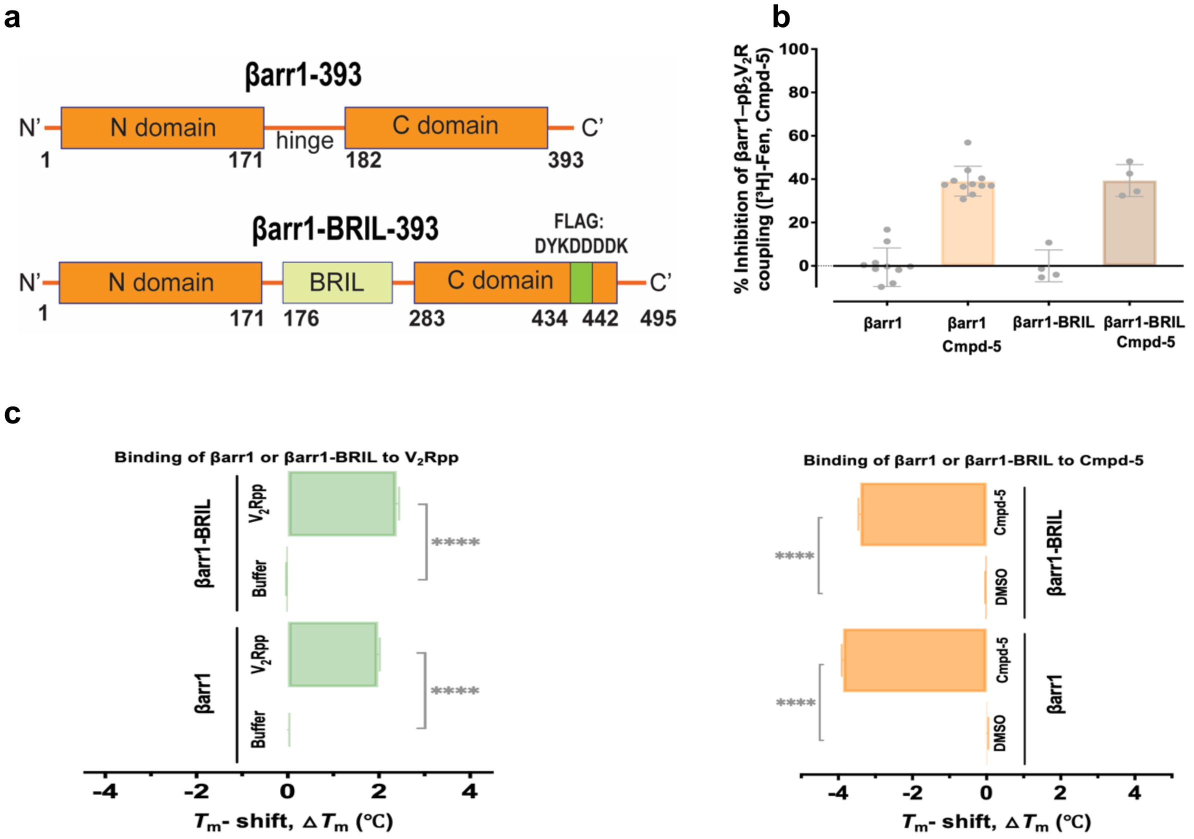


**Extended Data Fig. 6** | BRIL fusion to βarr1 preserves functional integrity for cryo-EM studies with Cmpd-5. **a**, Schematic of the βarr1–BRIL construct design. The BRIL domain was inserted into the hinge region of βarr1 as a fiducial marker for cryo-EM. Residue numbers in parentheses correspond to positions after the insertion of the apocytochrome b562 fusion protein (BRIL). **b**, Effect of Cmpd-5 (100 μM) on βarr1 (2 μM) and βarr1–BRIL (2 μM) promoted high-affinity agonist ([$^3$H]-Fenoterol, 6 nM) binding to $\beta_2V_2R$ *in vitro*. The data show that BRIL fusion does not impair the functional interaction between βarr1 and the receptor or Cmpd-5-mediated inhibition. Inhibition is expressed as a percentage relative to control. **c**, Binding of βarr1 and βarr1–BRIL to $V_2R$ phosphorylated peptide ($V_2$Rpp) (left) and Cmpd-5 (right), assessed by thermal stability shifts ($\Delta T_m$) using DSF. The results confirm that BRIL fusion does not alter βarr1 binding to either the $V_2$Rpp or Cmpd-5.

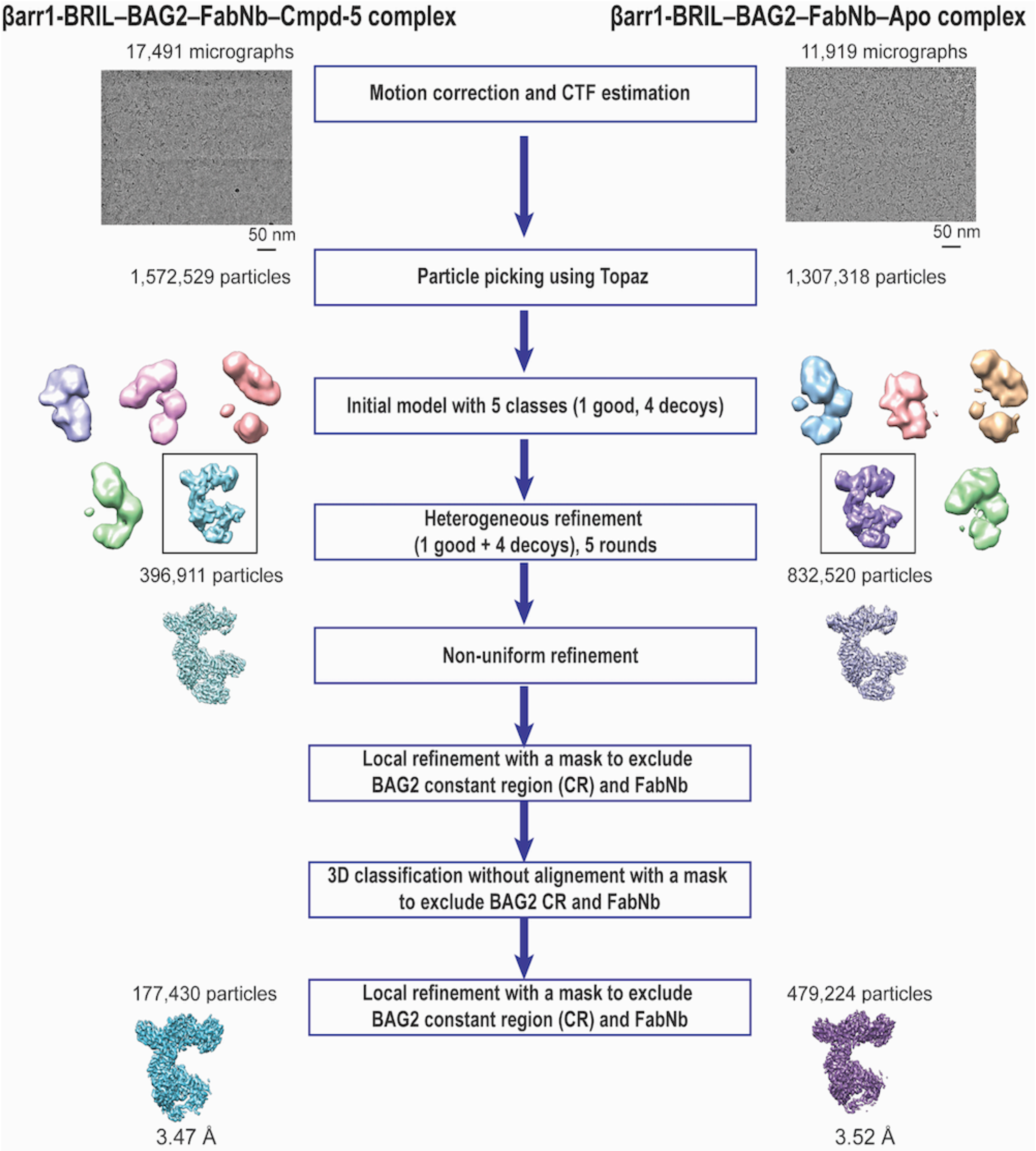


**Extended Data Fig. 7** | Flow chart of cryo-EM data processing for βarr1–BRIL–BAG2–aFabNb–Cmpd-5 and βarr1–BRIL–BAG2–aFabNb–Apo complexes.

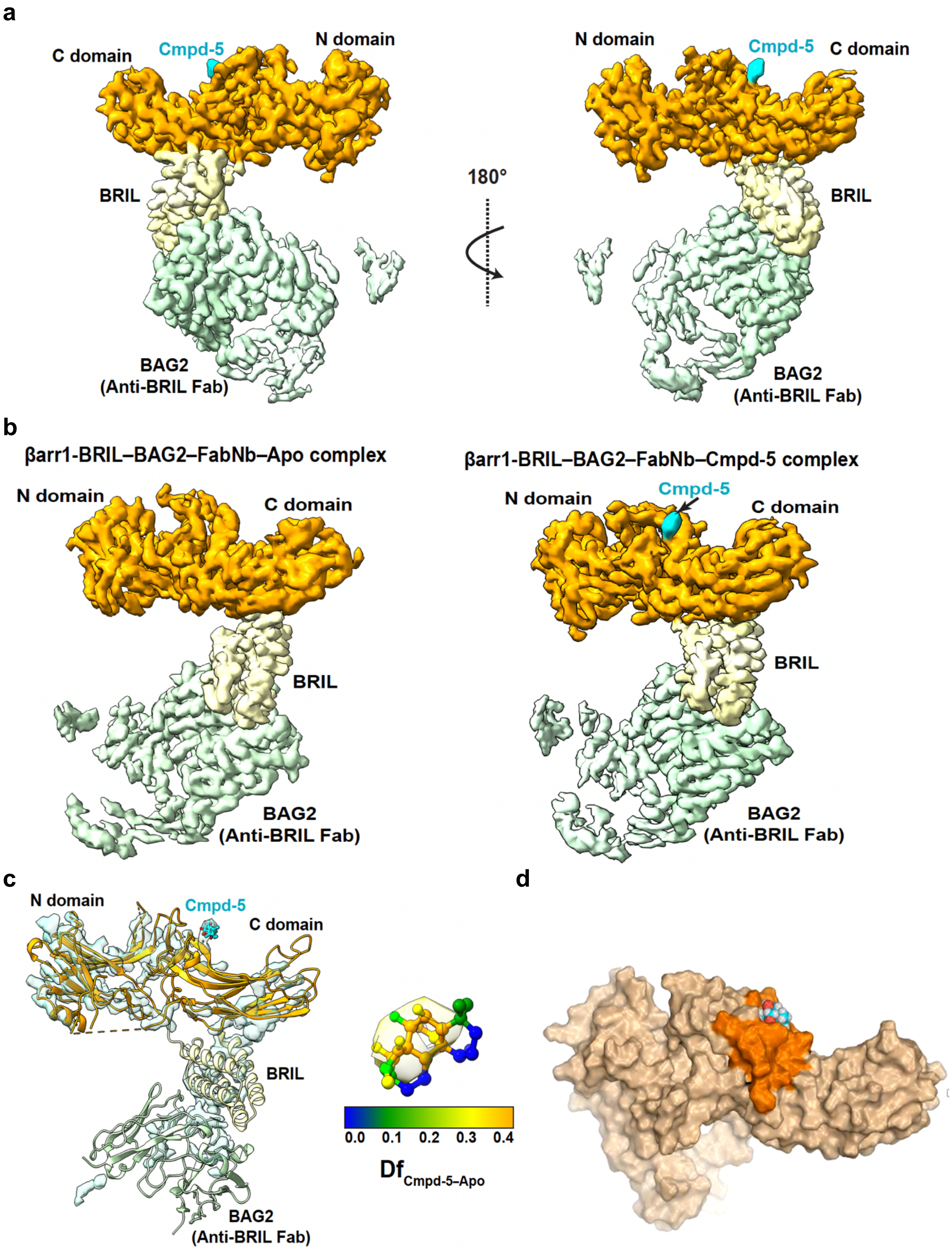
a
C domain
Cmpd-5
N domain
BRIL
BAG2
(Anti-BRIL Fab)
180°
N domain
Cmpd-5
C domain
BRIL
BAG2
(Anti-BRIL Fab)
b
βarr1-BRIL–BAG2–FabNb–Apo complex
N domain
C domain
BRIL
BAG2
(Anti-BRIL Fab)
βarr1-BRIL–BAG2–FabNb–Cmpd-5 complex
N domain
Cmpd-5
C domain
BRIL
BAG2
(Anti-BRIL Fab)
c
N domain
Cmpd-5
C domain
BRIL
BAG2
(Anti-BRIL Fab)
0.0 0.1 0.2 0.3 0.4
Df Cmpd-5–Apo
d

**Extended Data Fig. 8** | Cryo-EM density maps and structural insights into the βarr1–BRIL–BAG2–aFabNb–Cmpd-5 complex. **a,** Two views of the Cryo-EM density map of the βarr1–BRIL–BAG2–aFabNb–Cmpd-5 complex, colored by subunit (βarr1, green; BRIL, yellow; BAG2 (anti-BRIL Fab), blue; Cmpd-5, cyan). Post-processing was performed using DeepEMhancer[28] with a map contour level of 0.07. **b,** Cryo-EM density maps comparing the βarr1–BRIL–BAG2–aFabNb complex in the apo (3.52 Å, left) and Cmpd-5-bound (3.47 Å, right) states. Subunits are colored as in panel a. Maps were post-processed with DeepEMhancer[28] at a contour level of 0.07. **c**, Left, local density differences between the Cmpd-5-bound and apo states (difference map: Cmpd-5–Apo) calculated as described[43]. The atomic model of the Cmpd-5-bound state shows βarr1 (green), Cmpd-5 (cyan), BRIL (yellow), and BAG2 (blue). Right, density map of Cmpd-5 modeled with a ball-and-stick representation, colored by fractional density difference (Df, Cmpd-5–Apo) using a local scaling-based density difference approach. **d,** Molecular dynamics (MD) simulation snapshot showing Cmpd-5 binding within the MCL cleft of βarr1. The MCL cleft is colored orange, Cmpd-5 is cyan, and βarr1 is shown in wheat. This highlights the stable interaction of Cmpd-5 within the cleft.

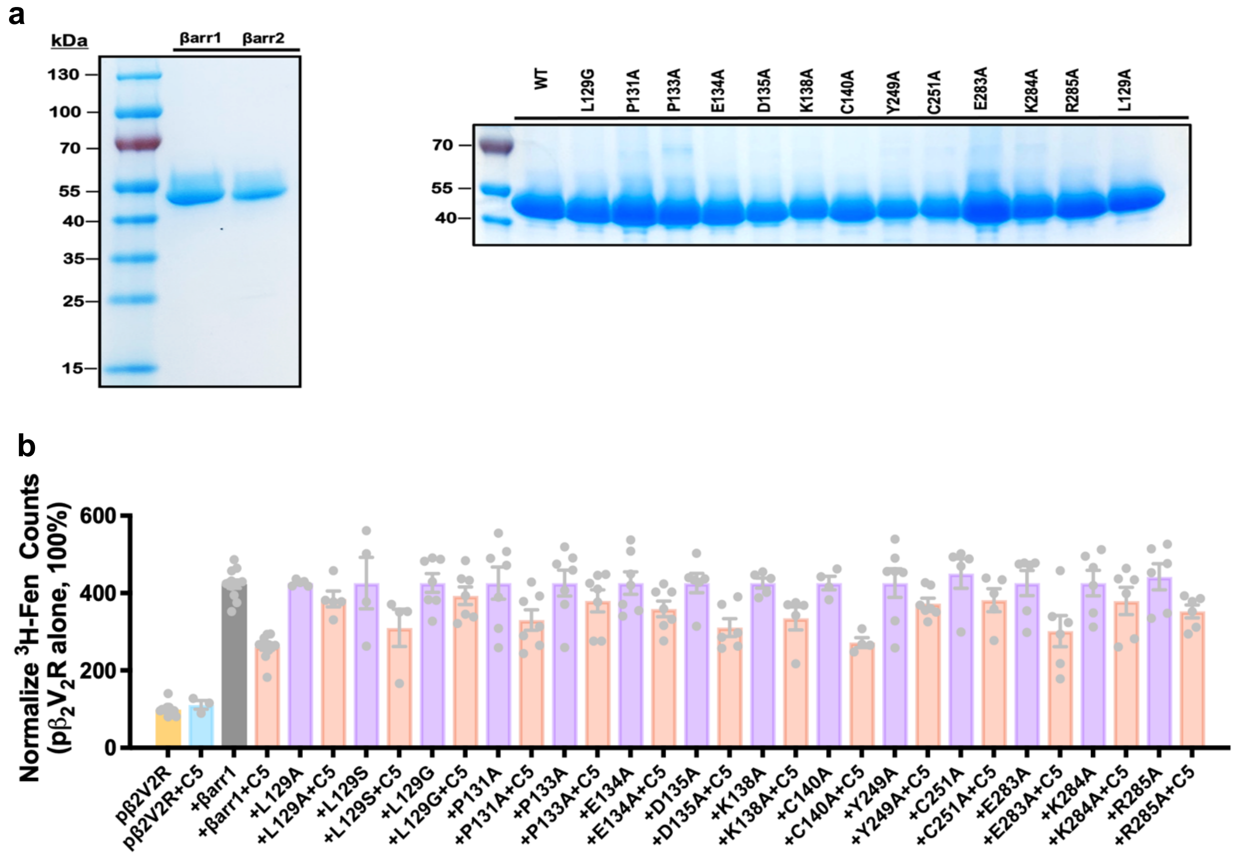

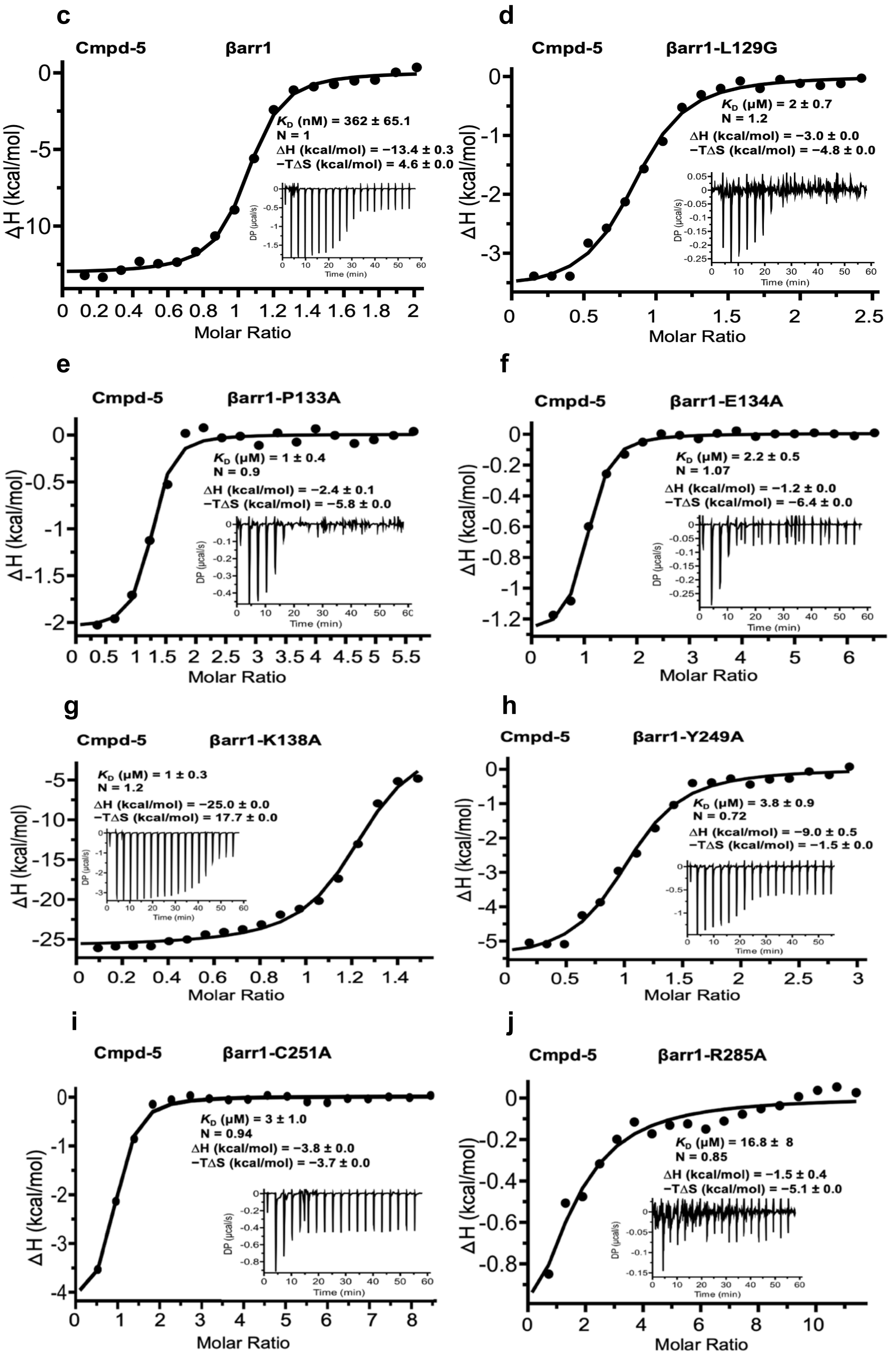
c
Cmpd-5 βarr1
$K_D$ (nM) = 362 ± 65.1
N = 1
ΔH (kcal/mol) = −13.4 ± 0.3
−TΔS (kcal/mol) = 4.6 ± 0.0
d
Cmpd-5 βarr1-L129G
$K_D$ (μM) = 2 ± 0.7
N = 1.2
ΔH (kcal/mol) = −3.0 ± 0.0
−TΔS (kcal/mol) = −4.8 ± 0.0
e
Cmpd-5 βarr1-P133A
$K_D$ (μM) = 1 ± 0.4
N = 0.9
ΔH (kcal/mol) = −2.4 ± 0.1
−TΔS (kcal/mol) = −5.8 ± 0.0
f
Cmpd-5 βarr1-E134A
$K_D$ (μM) = 2.2 ± 0.5
N = 1.07
ΔH (kcal/mol) = −1.2 ± 0.0
−TΔS (kcal/mol) = −6.4 ± 0.0
g
Cmpd-5 βarr1-K138A
$K_D$ (μM) = 1 ± 0.3
N = 1.2
ΔH (kcal/mol) = −25.0 ± 0.0
−TΔS (kcal/mol) = 17.7 ± 0.0
h
Cmpd-5 βarr1-Y249A
$K_D$ (μM) = 3.8 ± 0.9
N = 0.72
ΔH (kcal/mol) = −9.0 ± 0.5
−TΔS (kcal/mol) = −1.5 ± 0.0
i
Cmpd-5 βarr1-C251A
$K_D$ (μM) = 3 ± 1.0
N = 0.94
ΔH (kcal/mol) = −3.8 ± 0.0
−TΔS (kcal/mol) = −3.7 ± 0.0
j
Cmpd-5 βarr1-R285A
$K_D$ (μM) = 16.8 ± 8
N = 0.85
ΔH (kcal/mol) = −1.5 ± 0.4
−TΔS (kcal/mol) = −5.1 ± 0.0
ΔH (kcal/mol)
Molar Ratio
DP (μcal/s)
Time (min)

**Extended Data Fig. 9** | Mutagenesis, biochemical, functional, and biophysical validation of Cmpd-5 binding to βarr1. **a**, SDS–PAGE gels of purified recombinant βarr1–393, βarr2–394, and βarr1 point mutants targeting the Cmpd-5 binding pocket. Molecular weights were estimated using PageRuler™ Prestained Protein Ladder. **b**, Functional assay assessing Cmpd-5 inhibition of βarr1- and mutant-promoted high-affinity agonist binding to $β_2V_2R$ using [³H]-Fenoterol (6 nM) in Sf9 cell membranes. Data are normalized to p$β_2V_2R$ alone (100%) and represent mean ± s.e.m. from at least four independent experiments. **c–j**, ITC analysis of Cmpd-5 binding to βarr1 and its mutants at 25 °C. Representative thermograms (insets) and isotherms for (**c**) WT (also in Fig. 6E), (**d**) L129G, (**e**) P133A, (**f**) E134A, (**g**) K138A, (**h**) Y249A, (**i**) C251A, and (**j**) R285A. Data were fitted using a one-site binding model, reporting KD, stoichiometry (**n**), ΔH, and –TΔS with s.e.m.

**a**

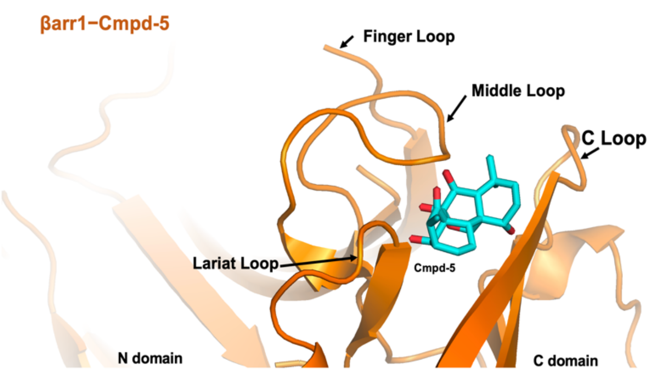


**b**

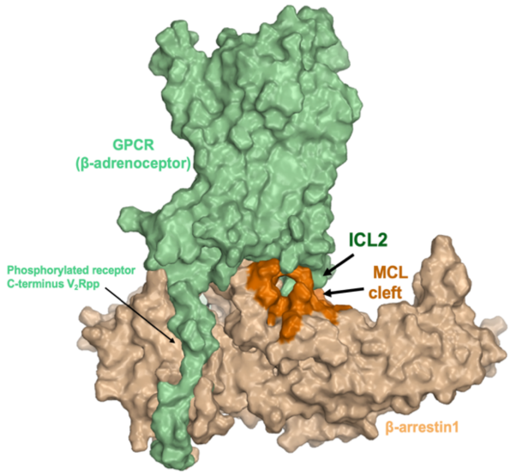


**c**

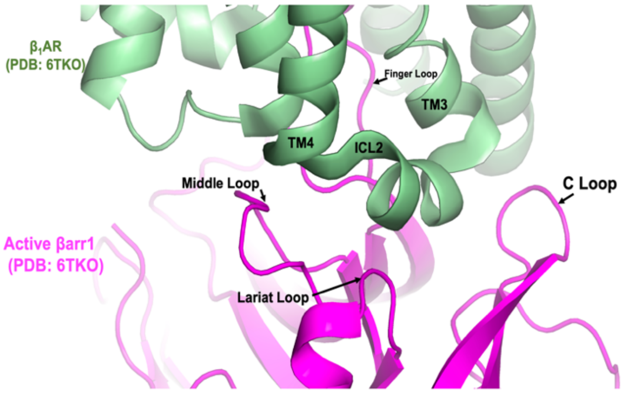


**d**

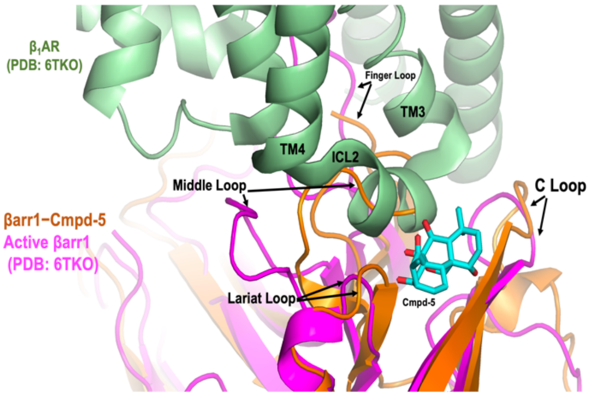

**e**

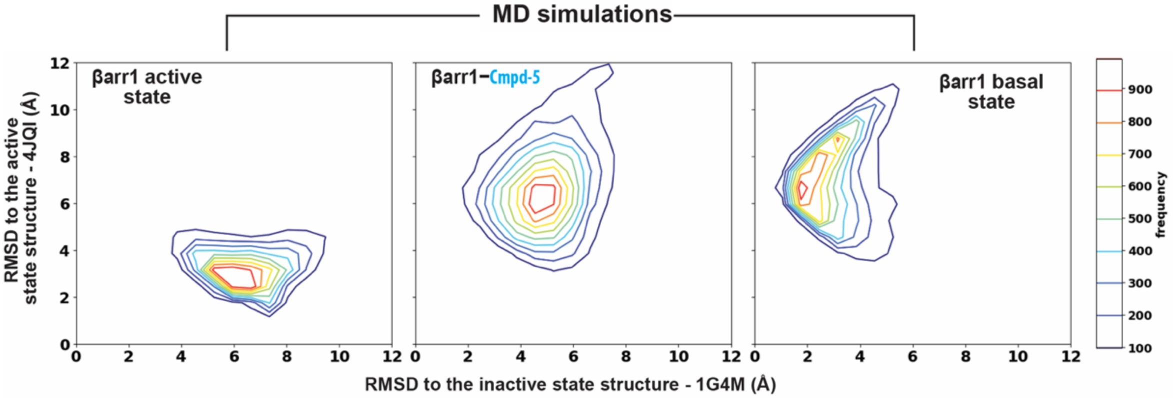


**Extended Data Fig. 10** | Structural basis and mechanism of Cmpd-5 binding and inhibition of βarr1. **a**–**d**, Structural basis of allosteric inhibition of βarr1 by Cmpd-5. **a**, Close-up view of the Cmpd-5 binding site within the central crest of βarr1 (orange), formed by the middle loop, C-loop, and lariat loop (MCL cleft). Cmpd-5 is shown in cyan. **b**, Molecular surface representation of the p$\beta_1V_2R$–βarr1 complex derived from cryo-EM structures (PDB: 6TKO). The p$\beta_1V_2R$ receptor is shown in pale green, βarr1 in wheat, and the MCL cleft in orange. Arrows indicate the MCL cleft, the ICL2 loop of GPCRs, and the phosphorylated C-terminal portion of p$\beta_1V_2R$ (pale yellow). **c**, Close-up view of the p$\beta_1V_2R$–βarr1 complex (PDB: 6TKO), with p$\beta_1V_2R$ in pale green and active βarr1 in magenta. **d**, Overlay of the cryo-EM structure of βarr1 bound to Cmpd-5 (cyan) with the $\beta_2V_2R$–βarr1 complex (PDB: 6TKO), highlighting conformational changes induced by Cmpd-5 binding. **e**, Cmpd-5 binding and conformational effects on βarr1 revealed by MD simulations. Two-dimensional (2D) contour plots showing the distribution of active, basal, and Cmpd-5-bound βarr1 states. The RMSD plot compares Cmpd-5-bound βarr1 from MD simulations relative to the basal (x-axis) and active (y-axis) states. The Cmpd-5–bound βarr1 adopts a conformation distinct from both canonical active and inactive states, preventing receptor engagement.

**Extended Data Table 1 |** Cryo-EM data collection, refinement, and validation statistics.

| | **βarr1–Cmpd-5 (βarr1-BRIL–BAG2-aFabNb–Cmpd-5)** (EMD-47042) (PDB 9DNM) | **βarr1 (Apo) (βarr1-BRIL–BAG2-aFabNb)** (EMD-47040) (PDB 9DNG) |
|---|---|---|
| **Data collection and processing** | | |
| Magnification | 81,000 | 81,000 |
| Voltage (kV) | 300 | 300 |
| Electron exposure (e–/Å$^2$) | 54.5 to 59.8 | 53.7 to 54.1 |
| Defocus range (μm) | -0.8 to -2.5 | -0.8 to -2.5 |
| Pixel size (Å) | 1.08 (collection) 1.3824 (final) | 1.08 (collection) 1.3824 (final) |
| Symmetry imposed | C1 | C1 |
| Initial particle projections (no.) | 1,572,529 | 1,307,318 |
| Final particle projections (no.) | 177,430 | 479,224 |
| Map resolution (Å) | 3.47 | 3.52 |
| FSC threshold | 0.143 | 0.143 |
| Map resolution range (Å) | 3.05-8.9 | 3.05-8.72 |
| **Refinement** | | |
| Initial model used (PDB code) | 1G4M | 1G4M |
| Model resolution (Å) | 3.4 | 3.5 |
| FSC threshold | 0.5 | 0.5 |
| Map sharpening *B* factor (Å$^2$) | -101.7 | -110.6 |
| Model composition | | |
| Non-hydrogen atoms | 5235 | 5252 |
| Protein residues | 668 | 675 |
| Ligands | 1 | 0 |
| *B* factors (Å$^2$) | | |
| Protein (min/max/mean) | 30.00/92.70/55.95 | 29.36/116.62/55.87 |
| Ligand | 77.01/77.01/77.01 | --- |
| R.m.s. deviations | | |
| Bond lengths (Å) | 0.005 | 0.004 |
| Bond angles (°) | 0.980 | 0.989 |
| Validation | | |
| MolProbity score | 1.76 | 1.81 |
| Clashscore | 7.96 | 7.46 |
| Poor rotamers (%) | 0.18 | 0.35 |
| Ramachandran plot | | |
| Favored (%) | 95.26 | 93.95 |
| Allowed (%) | 4.74 | 6.05 |
| Disallowed (%) | 0.00 | 0.00 |

**Extended Data Table 2 |** Summary of molecular dynamics simulations.

| state | number of trajectories | total length of simulations |
|---|---|---|
| Basal | 5 | 6600 |
| Cmpd-5-bound | 6 | 7200 |
| $V_2$Rpp-bound active | 5 | 7800 |

**Extended Data Table 3 |** Oligonucleotide primers for βarr1–Cmpd-5 site point mutations

| Oligonucleotides | |
|---|---|
| **Primer name** | **Sequence (5' to 3')** |
| βarr1-L129A_F | GCCCAGGTTGCGCTGTGACTGAGCACGGAAGGTT |
| βarr1-L129A_R | AACCTTCCGTGCTCAGTCACAGCGCAACCTGGGC |
| βarr1-L129S_F | GGCCCAGGTTGCGATGTGACTGAGCACG |
| βarr1-L129S_R | CGTGCTCAGTCACATCGCAACCTGGGCC |
| βarr1- L129G_F | GCCCAGGTTGCCCTGTGACTGAGCACGGAAGGTT |
| βarr1-L129G_R | AACCTTCCGTGCTCAGTCACAGGGCAACCTGGGC |
| βarr1-P131A_F | CTCAGGCCCAGCTTGCAATGTGACTGAGCAC |
| βarr1-P131A_R | GTGCTCAGTCACATTGCAAGCTGGGCCTGAG |
| βarr1-P133A_F | CCCTGTGTCCTCAGCCCCAGGTTGCAAT |
| βarr1-P133A_R | CATTGCAACCTGGGGCTGAGGACACAGGG- |
| βarr1-E134A_F | CTTCCCTGTGTCCGCAGGCCCAGGTTG |
| βarr1-E134A_R | CAACCTGGGCCTGCGGACACAGGGAAG |
| βarr1-D135A_F | GCCTTCCCTGTGGCCTCAGGCCCAG |
| βarr1-D135A_R | CTGGGCCTGAGGCCACAGGGAAGGC |
| βarr1-K138A_F | CCACACCGCAGGCCGCCCCTGTGTCCTCAG |
| βarr1-K138A_R | CTGAGGACACAGGGGCGGCCTGCGGTGTGG |
| βarr1-C140A_F | TTCATAATCCACACCGGCGGCCTTCCCTGTGTCC |
| βarr1-C140A_R | GGACACAGGGAAGGCCGCCGGTGTGGATTATGAA |
| βarr1-Y249A_F | GGCCACTGGGCACTTGGCCTGAGCTGTGTTGAAG |
| βarr1-Y249A_R | CTTCAACACAGCTCAGGCCAAGTGCCCAGTGGCC |
| βarr1-C251A_F | CTCCATGGCCACTGGGGCCTTGTACTGAGCTGTG |
| βarr1-C251A_R | CACAGCTCAGTACAAGGCCCCAGTGGCCATGGAG |
| βarr1-E283A_F | CAAGCCCCCGCTTCGCTCTGTTGTTTGCCA |
| βarr1-E283A_R | TGGCAAACAACAGAGCGAAGCGGGGGCTTG |
| βarr1-K284A_F | AGGGCAAGCCCCCGCGCCTCTCTGTTGTTTGC |
| βarr1-K284A_R | GCAAACAACAGAGAGGCGCGGGGGCTTGCCCT |
| βarr1-R285A_F | CGAGGGCAAGCCCCGCCTTCTCTCTGTTGT |
| βarr1-R285A_R | ACAACAGAGAGAAGGCGGGGCTTGCCCTCG |

**References**


1. Namkung, Y., *et al.* Monitoring G protein-coupled receptor and beta-arrestin trafficking in live cells using enhanced bystander BRET. *Nat Commun* **7**, 12178 (2016).
2. Smith, J.S., *et al.* Biased agonists of the chemokine receptor CXCR3 differentially control chemotaxis and inflammation. *Sci Signal* **11**(2018).
3. Rein, L.A., *et al.* beta-Arrestin2 mediates progression of murine primary myelofibrosis. *JCI Insight* **2**(2017).
4. Shukla, A.K., *et al.* Visualization of arrestin recruitment by a G-protein-coupled receptor. *Nature* **512**, 218-222 (2014).
5. Shukla, A.K., *et al.* Structure of active beta-arrestin-1 bound to a G-protein-coupled receptor phosphopeptide. *Nature* **497**, 137-141 (2013).
6. Cahill, T.J., 3rd, *et al.* Distinct conformations of GPCR-beta-arrestin complexes mediate desensitization, signaling, and endocytosis. *Proc Natl Acad Sci U S A* **114**, 2562-2567 (2017).
7. Nobles, K.N., Guan, Z., Xiao, K., Oas, T.G. & Lefkowitz, R.J. The active conformation of beta-arrestin1: direct evidence for the phosphate sensor in the N-domain and conformational differences in the active states of beta-arrestins1 and -2. *J Biol Chem* **282**, 21370-21381 (2007).
8. Rasmussen, S.G., *et al.* Crystal structure of the beta2 adrenergic receptor-Gs protein complex. *Nature* **477**, 549-555 (2011).
9. Kahsai, A.W., *et al.* Signal transduction at GPCRs: Allosteric activation of the ERK MAPK by beta-arrestin. *Proc Natl Acad Sci U S A* **120**, e2303794120 (2023).
10. Pakharukova, N., Masoudi, A., Pani, B., Staus, D.P. & Lefkowitz, R.J. Allosteric activation of proto-oncogene kinase Src by GPCR-beta-arrestin complexes. *J Biol Chem* **295**, 16773-16784 (2020).
11. Wilkins, M.R., *et al.* Protein identification and analysis tools in the ExPASy server. *Methods Mol Biol* **112**, 531-552 (1999).
12. Kahsai, A.W., *et al.* Conformationally selective RNA aptamers allosterically modulate the beta2-adrenoceptor. *Nat Chem Biol* **12**, 709-716 (2016).
13. Kahsai, A.W., *et al.* Multiple ligand-specific conformations of the beta2-adrenergic receptor. *Nat Chem Biol* **7**, 692-700 (2011).
14. Rasmussen, S.G., *et al.* Structure of a nanobody-stabilized active state of the beta(2) adrenoceptor. *Nature* **469**, 175-180 (2011).
15. Ahn, S., *et al.* Allosteric "beta-blocker" isolated from a DNA-encoded small molecule library. *Proc Natl Acad Sci U S A* **114**, 1708-1713 (2017).
16. Dixon, A.S., *et al.* NanoLuc Complementation Reporter Optimized for Accurate Measurement of Protein Interactions in Cells. *ACS Chem Biol* **11**, 400-408 (2016).
17. Daly, C., *et al.* beta-Arrestin-dependent and -independent endosomal G protein activation by the vasopressin type 2 receptor. *Elife* **12**(2023).
18. Pandey, S., *et al.* Intrinsic bias at non-canonical, beta-arrestin-coupled seven transmembrane receptors. *Mol Cell* **81**, 4605-4621 e4611 (2021).

19. Violin, J.D., *et al.* beta2-adrenergic receptor signaling and desensitization elucidated by quantitative modeling of real time cAMP dynamics. *J Biol Chem* **283**, 2949-2961 (2008).
20. Li, A., Liu, S., Huang, R., Ahn, S. & Lefkowitz, R.J. Loss of biased signaling at a G protein-coupled receptor in overexpressed systems. *PLoS One* **18**, e0283477 (2023).
21. Smith, J.S., *et al.* C-X-C Motif Chemokine Receptor 3 Splice Variants Differentially Activate Beta-Arrestins to Regulate Downstream Signaling Pathways. *Mol Pharmacol* **92**, 136-150 (2017).
22. Wisler, J.W., *et al.* A unique mechanism of beta-blocker action: carvedilol stimulates beta-arrestin signaling. *Proc Natl Acad Sci U S A* **104**, 16657-16662 (2007).
23. Wang, J., *et al.* Galphai is required for carvedilol-induced beta1 adrenergic receptor beta-arrestin biased signaling. *Nat Commun* **8**, 1706 (2017).
24. Mukherjee, S., *et al.* Synthetic antibodies against BRIL as universal fiducial marks for single-particle cryoEM structure determination of membrane proteins. *Nat Commun* **11**, 1598 (2020).
25. Ereno-Orbea, J., *et al.* Structural Basis of Enhanced Crystallizability Induced by a Molecular Chaperone for Antibody Antigen-Binding Fragments. *J Mol Biol* **430**, 322-336 (2018).
26. Punjani, A., Rubinstein, J.L., Fleet, D.J. & Brubaker, M.A. cryoSPARC: algorithms for rapid unsupervised cryo-EM structure determination. *Nat Methods* **14**, 290-296 (2017).
27. Bepler, T., *et al.* Positive-unlabeled convolutional neural networks for particle picking in cryo-electron micrographs. *Nat Methods* **16**, 1153-1160 (2019).
28. Sanchez-Garcia, R., *et al.* DeepEMhancer: a deep learning solution for cryo-EM volume post-processing. *Commun Biol* **4**, 874 (2021).
29. Han, M., Gurevich, V.V., Vishnivetskiy, S.A., Sigler, P.B. & Schubert, C. Crystal structure of beta-arrestin at 1.9 A: possible mechanism of receptor binding and membrane Translocation. *Structure* **9**, 869-880 (2001).
30. Pettersen, E.F., *et al.* UCSF Chimera--a visualization system for exploratory research and analysis. *J Comput Chem* **25**, 1605-1612 (2004).
31. Tsutsumi, N., *et al.* Structure of human Frizzled5 by fiducial-assisted cryo-EM supports a heterodimeric mechanism of canonical Wnt signaling. *Elife* **9**(2020).
32. Emsley, P., Lohkamp, B., Scott, W.G. & Cowtan, K. Features and development of Coot. *Acta Crystallogr D Biol Crystallogr* **66**, 486-501 (2010).
33. Croll, T.I. ISOLDE: a physically realistic environment for model building into low-resolution electron-density maps. *Acta Crystallogr D Struct Biol* **74**, 519-530 (2018).
34. Adams, P.D., *et al.* The Phenix software for automated determination of macromolecular structures. *Methods* **55**, 94-106 (2011).
35. Chen, V.B., *et al.* MolProbity: all-atom structure validation for macromolecular crystallography. *Acta Crystallogr D Biol Crystallogr* **66**, 12-21 (2010).
36. Lee, Y., *et al.* Molecular basis of beta-arrestin coupling to formoterol-bound beta(1)-adrenoceptor. *Nature* **583**, 862-866 (2020).

37. Asher, W.B., *et al.* GPCR-mediated beta-arrestin activation deconvoluted with single-molecule precision. *Cell* **185**, 1661-1675 e1616 (2022).
38. Webb, B. & Sali, A. Comparative Protein Structure Modeling Using MODELLER. *Curr Protoc Bioinformatics* **54**, 5 6 1-5 6 37 (2016).
39. Bowers, K.J., *et al.* Scalable Algorithms for Molecular Dynamics Simulations on Commodity Clusters. in *ACM/IEEE SC 2006 Conference (SC'06)* 43 (2006).
40. Lu, C., *et al.* OPLS4: Improving Force Field Accuracy on Challenging Regimes of Chemical Space. *J Chem Theory Comput* **17**, 4291-4300 (2021).
41. Feller, S.E., Zhang, Y.H., Pastor, R.W. & Brooks, B.R. Constant-Pressure Molecular-Dynamics Simulation - the Langevin Piston Method. *J Chem Phys* **103**, 4613-4621 (1995).
42. Humphrey, W., Dalke, A. & Schulten, K. VMD: visual molecular dynamics. *J Mol Graph* **14**, 33-38, 27-38 (1996).
43. Joseph, A.P., *et al.* Comparing Cryo-EM Reconstructions and Validating Atomic Model Fit Using Difference Maps. *J Chem Inf Model* **60**, 2552-2560 (2020).
